# Auditory corticofugal neurons transmit auditory and non-auditory information during behavior

**DOI:** 10.1101/2022.08.08.503214

**Authors:** Alexander N. Ford, Jordyn E. Czarny, Meike M. Rogalla, Gunnar L. Quass, Pierre F. Apostolides

## Abstract

Layer 5 pyramidal neurons of sensory cortices project “corticofugal” axons to myriad sub-cortical targets, thereby broadcasting high-level signals important for perception and learning. Recent studies suggest *dendritic Ca^2+^ spikes* as key biophysical mechanisms supporting corticofugal neuron function: These long-lasting events drive burst firing, thereby initiating uniquely powerful signals to modulate sub-cortical representations and trigger learning-related plasticity. However, the behavioral relevance of corticofugal dendritic spikes is poorly understood. We shed light on this issue using 2-photon Ca^2+^ imaging of auditory corticofugal dendrites as mice of either sex engage in a GO/NO-GO sound-discrimination task.

Unexpectedly, only a minority of dendritic spikes were triggered by behaviorally relevant sounds under our conditions. Task related dendritic activity instead mostly followed sound cue termination and co-occurred with mice’s instrumental licking during the answer period of behavioral trials, irrespective of reward consumption. Temporally selective, optogenetic silencing of corticofugal neurons during the trial answer period impaired auditory discrimination learning. Thus, auditory corticofugal systems’ contribution to learning and plasticity may be partially non-sensory in nature.

**Significance Statement:** The auditory cortex sends a massive “feedback” projection to the inferior colliculus (IC) which controls IC neuron plasticity and some types of perceptual learning. Precisely what signals are fed back during behavior is unclear. Using multiphoton imaging of auditory cortico-collicular neurons as mice engage in a sound discrimination task, we find that activity coincides more with mice’s instrumental actions rather than sound cues. Dendritic Ca^2+^ spikes and burst firing contributed to this non-auditory activity, which is notable given that dendritic spikes instruct synaptic plasticity in many other circuits. Accordingly, optogenetic silencing of corticofugal neurons during mice’s instrumental actions impaired discriminative learning. Auditory corticofugal neurons may thus transmit significant non-auditory information that contributes to learning-related plasticity.

## Introduction

A ubiquitous circuit motif of the neo-cortex is that layer 5 “corticofugal” pyramidal neurons send axons to multiple sub-cortical regions, thereby broadcasting cortical computations throughout the brain (Usrey and Sherman, 2019; Sherman and Usrey, 2021). Consequently, these descending pathways enable a predictive or feedback control of hierarchically lower circuits that influences learning and behavior across multiple time scales (Bajo et al., 2010; Liu et al., 2016; Takahashi et al., 2016, 2020; Ranganathan et al., 2018; Doron et al., 2020; Ruediger and Scanziani, 2020). Interestingly, corticofugal neurons generate two distinct types of action potential patterns: Low frequency (1-50 Hz) simple spike trains, and powerful high frequency bursts (100-200 Hz) driven by a long-lasting, regenerative Ca^2+^ spike initiated in the apical dendrites (Larkum et al., 1999; Harnett et al., 2013; Slater et al., 2013; Beaulieu-Laroche et al., 2019; Fletcher and Williams, 2019; Lacefield et al., 2019; Galloni et al., 2020). Because the neo-cortex is suggested to transmit information via a rate code (Banerjee et al., 2008; London et al., 2010), burst firing may be a highly salient output signal with important downstream consequences. Indeed, compared to low-frequency spike trains, bursts often result in greater temporal summation (Lisman, 1997; Tsodyks and Markram, 1997; Kim and McCormick, 1998; Apostolides et al., 2016), synaptic facilitation (Kreitzer and Regehr, 2000; Xu et al., 2012b), post-tetanic potentiation (Williams and Atkinson, 2007; Neubrandt et al., 2018), and extrasynaptic spillover (Clark and Cull-Candy, 2002; Hires et al., 2008; Nahir and Jahr, 2013) at downstream targets. Bursts could thus multiplex unique signals that function synergistically with simple spike trains, thereby increasing the computational flexibility of cortical circuits (Naud and Sprekeler, 2018; Payeur et al., 2021). However, the behavioral relevance of dendritic spikes is unclear, owing to the difficulty of recording dendritic activity in actively engaged animals.

We address this knowledge gap using 2-photon Ca^2+^ imaging of layer 5 corticofugal neuron dendrites in auditory cortex of mice performing a reward-based, GO/NO-GO sound discrimination task. We used a retrograde adeno-associated virus (AAV; Tervo et al., 2016) to express the Ca^2+^ indicator GCaMP6f in auditory corticofugal neurons projecting to the inferior colliculus (IC), a major midbrain hub that integrates ascending auditory brainstem signals with descending feedback from auditory cortex (Morest and Oliver, 1984; Winer et al., 1998; Oberle et al., 2022). Interestingly many auditory corticofugal dendrites were *inhibited* by sound, whereas a subset of neurons generated dendritic spikes preferentially in response to sounds predicting reward availability. Rather, a major fraction of task-related dendritic activity followed sound termination and occurred synchronously with mice’s operant licking during the trial answer period, independent of reward consumption. Optogenetic silencing of corticofugal neurons selectively during the trial answer period impaired discriminative auditory learning, suggesting a non-sensory contribution of auditory cortex to learned behaviors. The data more broadly imply that non-auditory signals recruit plasticity mechanisms, either in corticofugal neurons or their downstream targets, to stamp in learned associations between sensations and instrumental actions.

## Methods

### Surgeries

All surgical procedures were approved by the University of Michigan’s IACUC, performed in accordance with NIH’s guide for the care and use of laboratory animals, and conducted under aseptic conditions. C57BL6/J mice from Jackson labs (5-7 weeks of age at time of surgery; stock #000664) were used for most imaging experiments. In experiments of Figure 8, we used F1 offsprings of C57BL6/J x CBA/CaJ matings born in our colony (6-8 weeks of age at time of surgery). Mice were deeply anesthetized with 4-5% isoflurane vaporized in O_2_ and mounted in a rotating stereotax (David Kopf Instruments). Isoflurane was subsequently lowered to 1-2% to maintain a stable anesthetic plane, as assessed by the absence of paw withdrawal reflex and stable respiration (1-1.5 breaths/s). Body temperature was maintained near 37-38 C using a feedback controlled, homeothermic heating blanket (Harvard Apparatus). Mice were administered carprofen (5 mg/kg, s.c.) as a pre-surgical analgesic, the fur and skin overlaying the skull was removed and 2% topical lidocaine was applied to the wound margins. The dorsal portion of the skull was cleaned and scored in a checkerboard pattern using a scalpel. A 300-500 µm craniotomy was carefully drilled over the left IC (0.9 mm caudal and 1.0 mm lateral from the lamboid sutures) using a 0.5 mm dental burr and microdrill (Foredom). The skull was frequently irrigated with chilled PBS to prevent heating, and care was taken as to not rupture the transverse sinus or overlaying dura mater during drilling. Following the craniotomy, a glass pipette (tip diameter: ∼100 µm) containing the retrograde AAV-hSyn1-GCaMP6f-P2A-nls-dTomato virus (Addgene #51085) was slowly lowered into the IC at a rate of 1 µm/s using a micromanipulator (Sutter MP-285). In N = 2 mice used in Figure 8, we injected a retrograde AAV-syn-jGCaMP8s-WPRE (Addgene #162374) in place of the GCaMP6f virus listed above. A total of 125-165 nL was injected at 5 different sites in the IC (25-33 nL/site, 1 nL/s) per mouse. Following injections, the pipette was maintained in place for an additional 3-5 min before retraction at a rate of 10 µm/s. The craniotomy over the IC was then filled with bone wax and the dorsal surface of the skull was covered with a thin layer of cyanoacrylate glue.

Subsequently, the left temporal muscle was retracted and the stereotax was rotated ∼50 degrees, thereby enabling a vertical approach to the lateral left portion of the mouse’s skull. A ∼2 mm diameter craniotomy was carefully opened over the left auditory cortex (2.75 mm caudal from bregma centered over the lateral ridge). The dura overlying the auditory cortex was left intact, and a custom cranial window (two 2 mm diameter coverslips stacked upon one another and affixed to a 4 mm outer coverslip with UV-cured Norland Adhesive #71) was gently implanted over the exposed brain. The coverslip was affixed to the skull using cyanoacrylate glue and reinforced with dental cement (Lang dental). Following cranial window implantation, the rotating stereotax was repositioned horizontally and a lightweight titanium headbar was affixed to the mouse’s skull using dental cement. The mouse was then removed from the stereotax, administered a post-operative analgesic (buprenorphine, 0.03 mg/kg) and allowed to recover from surgery in a clean home cage on a heating pad. Mice were then returned to the vivarium. Supplemental doses of carprofen were administered 24 and 48 hours following surgery.

For behavior and optogenetic experiments of Figure 12-13, surgeries were conducted as described above, except that a retrograde-cre virus (pENN.AAV.hSyn.Cre.WPRE.hGH; Addgene #105553) or sterile PBS was injected bilaterally in the IC of 6-8 week old GtACR1 fl/fl mice (Jackson stock # 033089) and 4 mm diameter coverslips were implanted over the thinned skull of both auditory cortices. 4-5 week old GtACR1 mice were used for the experiments of Figure 9. Here, an incision was gently made in the skin overlying the skull, a hole was drilled over the left IC to unilaterally inject ∼100 nL of retrograde-cre virus. Following injections, the pipette was retracted, the skin was sutured and post-operative care was provided as described above.

### Behavior during 2-photon Ca^2+^ Imaging

Mice were single housed under a reversed light/dark cycle (12/12 hr). Food, and initially water, were provided ad libitum. 3-7 days following surgery, mice’s water ration was restricted to 0.5-1.5 mL water/day, depending on weight loss and motivation. Body weight was monitored daily and kept above 70 % of the ad libitum weight. Following 5-7 days of water restriction, mice were handled by an experimenter and habituated to head-fixation while sitting in a Plexiglas tube, similar to the protocol described by Guo and colleagues (2014). After one to two initial habituation sessions of ∼30 min each, mice were trained on the auditory GO/NO-GO discrimination task once per day in custom-made sound-attenuating booths. The behavioral protocol was controlled by BPod hardware and related Matlab code (Sanworks). During behavioral sessions, an optical lickport (Sanworks) connected to a reward spout was placed within 1-2 cm from the mouse’s mouth. On each trial, a 1-s sound (band-passed white noise, 4-16 kHz bandwidth; 0 or 100% 15 Hz modulation depth, calibrated to 70 dB with a ¼” microphone [Bruel & Kjaer]) was presented through a speaker (Peerless XT25SC90-04) located horizontally ∼10 cm away from the mouse’s right ear. Stimulus category (GO or NO-GO) was pseudo randomly assigned as 0 or 100 % sAM depth for each animal. On GO trials, licking the reward spout during a 1.5 s “answer period” following sound termination resulted in the opening of a solenoid valve (located outside of the sound attenuating chamber) and delivery of a ∼ 2 µL drop of 10% sucrose water. On NO-GO trials, licking the water spout during the answer period resulted in a “time-out”, where the inter-trial interval increased by 5-10 s. Licking the water spout at any other time during the trial had no effect, and consequently mice learned to lick mainly during the answer period on GO trials after 5-10 training sessions. Behavioral sessions consisted of approximately equal numbers of randomly presented GO and NO-GO trials, with no more than three consecutive trials being of the same trial type. Sessions typically lasted 1.5-3 hours, during which time mice performed ∼250-600 trials. All mouse training and data collection were conducted during mice’s dark cycle.

Imaging data acquisition typically commenced once mice performed at a global average of 75-80% correct for a minimum of two consecutive training sessions. Mice underwent a maximum of 5 imaging sessions/week (maximum one per day, Monday-Friday) for 2 to 5 weeks following surgery. At the beginning of each imaging session, a temporary silicone well was built around the cranial window to hold saline solution. Experiments were performed using a resonant-galvo 2-photon microscope similar to the Janelia MIMMs design (Sutter Instruments) and a water immersion 16x objective (0.8 NA; Nikon N16XLWD-PF). The microscope was housed in a custom sound-attenuating chamber. The microscope objective was rotated 40-50 degrees to be approximately orthogonal to the cranial window over auditory cortex. GCaMP6f was excited with a Titanium-Sapphire laser at 920 nm (Coherent Chameleon Ultra II). During each session, we imaged a single field of view (∼350 µm^2^) 150-300 µm below the pial surface containing apical trunk and/or tuft dendrites of layer 5 cortico-collicular pyramidal neurons. Although our data were collected without explicitly defining auditory cortical sub-fields, layer 5 auditory cortico-collicular neurons are mostly restricted to the primary auditory cortex and anterior auditory field (Yudintsev et al., 2021). Typically, data could be acquired from 2 to 8 non-overlapping fields of view per mouse. On each trial, 7-8 s of imaging data were acquired with Scanimage software at 30 Hz, with a minimum trial-to-trial interval of 8 s. Behavior data were acquired simultaneously and synchronized with imaging data offline. At the end of each session, the silicone well was removed and mice were returned to their home cage. Mice’s water intake was estimated by calculating the difference in body weight (including droppings) before and after each behavior session. Mice were provided with additional water (0% sucrose) in their home cage if they did not receive their full daily water ration during the sessions. Similarly, mice were provided with appropriate rations of 0% sucrose water on days in which they did not participate in behavioral experiments.

### Behavior during optogenetic manipulations

Surgeons and experimenters were blinded to mice’s treatment condition throughout the course of these experiments. Following a week of recovery from surgery, mice were placed on water restriction and handled by the experimenter for 7 days prior to training. Three days prior to training, all mice were habituated to head fixation for sessions of up to 30 min over a three day period. All mice followed a standardized training protocol whereby head-fixed mice were first trained on 250 trials/day to lick a water-spout within the answer period (2 s duration for all optogenetic experiments) following a GO cue to receive a drop of sugar water reward (band-pass white noise, 4-16 kHz, 1 s duration, 75-82 dB SPL for all mice). Upon reaching criterion of >75% hits/session for 2 consecutive sessions, the NO-GO cue (4-16 kHz band-pass white noise with 15 Hz modulation rate, 75-82 dB SPL) was introduced in the following session and false alarms were punished with a 7 s timeout as described above. In the first session following criterion, mice initially received 100 sequential GO trials, followed by a total of 200 interleaved GO and NO-GO trials (50% NO-GO probability). This initial “transition session” was discarded from analyses. All subsequent sessions contained 300 pseudo-randomly interleaved GO and NO-GO trials (50% NO-GO probability), with no more than 3 sequential repetitions of the same trial type. Auditory corticofugal neurons were silenced for 2 s during the answer period via 625 nm LED coupled optic fibers (400 µm diameter, 0.5 NA) positioned atop cranial windows overlaying the auditory cortices. LED power at the fiber tip was 10-12 mW. We estimated the power loss through the skull and brain to be ∼1 mW. To this end, we compared the power output of the 625 nm LED, before and after placing a mouse parietal skull plate lined with ∼1 mm thick 2% agarose between the optic fiber and the power meter detector. We used 625 instead of 473 nm light due to the decreased scattering of longer wavelengths in brain tissue. In addition, prior experiments in neo-cortex (Li et al., 2019) and our in vitro calibration data (Figure 9) show robust GtACR1 activation at 625 nm.

### Imaging data analysis

Movies were registered and motion corrected with the Suite2p package running in Matlab or Python (Pachitariu et al., 2016). ROIs for individual dendrite segments were generated using the Caiman package running in Matlab 2017b (Pnevmatikakis et al., 2016; Lacefield et al., 2019). Fluorescence time series of individual ROIs were generated by averaging the frame-by-frame intensity of pixels corresponding to binary masks of dendrite segments. To convert raw fluorescence to ΔF/F, baseline fluorescence of each ROI was defined as the median pixel intensity over a sliding window of 20-30 trials (160-240 s of data).

Spatially non-overlapping ROIs in the same FOV occasionally showed high degrees of temporal correlation, in agreement with previous studies showing that dendritic Ca^2+^ transients in layer 5 pyramidal neurons are often globally synchronized across multiple dendritic branches (Xu et al., 2012a; Ranganathan et al., 2018; Beaulieu-Laroche et al., 2019; Kerlin et al., 2019). We considered highly correlated dendrites (>0.5 cross-correlation coefficient; Ranganathan et al., 2018) as originating from the same neuron, and thus only a single ROI was kept for the analyses reported here. The timing and peak of Ca^2+^ events were detected using a threshold crossing algorithm (peakfinder, Nathanael Yoder, Matlab Central File Exchange #25500), after smoothing traces with a 4-point median filter. Thresholds were set between 0.5 and 1.0 ΔF/F depending on the ROI. Time series and detected events were visually inspected and curated to exclude occasional false positives. We did not apply neuropil subtraction to the fluorescence signals because our retrograde labeling approach resulted in sparse GCaMP6f expression and generally low baseline fluorescence values. Neuropil subtraction under these conditions risks over-correcting the fluorescence time series, resulting in artifactual ΔF/F signals.

Normalized event rate histograms in Figure 10 were calculated as follows: For each task modulated dendrite, we quantified the probability of observing a dendritic event at each time bin (33 ms bin width) across each trial. We then divided the value of each bin by the mean event probability observed during a 2-3 s “baseline” period prior to sound onset. To test for spatial clustering of dendritic response profiles (Figure 4), we first calculated the centroid of each ROI’s binary mask using the Matlab function regionprops(). We then calculated the pairwise Euclidean distance between all task responsive ROIs in a given FOV.

### In vitro electrophysiology

∼2 weeks following surgery, mice were deeply anesthetized with isoflurane, decapitated, and the brains quickly removed in ice cold, oxygenated solution containing (in mM): 87 NaCl, 2.5 KCl, 1.25 NaH_2_PO_4_, 7 MgCl_2_, 25 glucose, 75 sucrose, 25 NaHCO_3_, 1 ascorbate, 1 pyruvate, 0.5 CaCl_2_. 300 μm thick coronal slices containing the auditory cortex were sectioned with a vibratome (Campden instruments) followed by a 30 min recovery incubation in oxygenated, 34 C artificial cerebro-spinal fluid (ACSF; in mM): 119 NaCl, 25 NaHCO_3_, 3 KCl, 1.25 NaH_2_PO_4_, 15 glucose, 1 MgCl_2_, 1.3 CaCl_2_, 1 ascorbate, 3 pyruvate. Following recovery, a slice was transferred to a recording chamber under an Olympus BXW51 microscope and continuously perfused with oxygenated 32-34° C ACSF (2-4 mL/min, chamber volume: ∼1 mL). Neurons were visualized via Dodt contrast optics using a 63x microscope objective and targeted for patch-clamp recordings based on visually confirmed expression of the FusionRed tag fused to the GtACR1 construct (Li et al., 2019). Current-clamp recordings were obtained with glass pipettes filled with internal solution (in mM): 115 K-gluconate, 4 KCl, 0.1 EGTA, 10 HEPES, 14 Tris-phosphocreatine, 4 Mg-ATP, 0.5 Tris-GTP, 4 NaCl, pH 7.2-7.3, 290 mOsm (open tip resistance: 3-5 MOhm). 91% of neurons recorded for these experiments had onset doublets or burst firing patterns characteristics of thick-tufted, pyramidal tract-type Layer 5 neurons (10/11 cells; 8/9 cells used for GtACR1 calibration in Figure 11, and 2/2 cells excluded from the analyses due to incomplete data collection). These observations are in qualitative agreement with the reported physiology of auditory cortico-collicular neurons (Slater et al., 2013; Joshi et al., 2015). GtACR1 was activated via a 400 µm diameter optic fiber coupled to a 625 nM LED positioned <1 mm from the recorded cell. Control and LED trials were interleaved during experiments, and we typically recorded 3-4 repetitions of each current step/LED power combination. Data were acquired with an Axon Instruments Multiclamp 700B, low-pass filtered online at 10 kHz, and sampled at 50 kHz with a National instruments PCIe-6343 card and BNC2090A interface controlled by Matlab based Wavesurfer software (Janelia Research Campus). Series resistance (<25 mOhm) was compensated via bridge balance circuitry and pipette capacitance neutralization was employed in all experiments.

### Statistics

All statistical routines were run in Matlab, Graphpad Prism 9, or Microsoft Excel. The d-prime was calculated as the difference in z-statistic between hit and false alarm rates across all viable trials in a given session. All analyses were run on task-modulated dendritic ROIs, which were defined as having a Ca^2+^ event probability during the sound and/or answer periods that was significantly different from the baseline period on either GO or NO-GO trials (two-tailed Wilcoxon signed-rank test). P-values were adjusted for 6 multiple comparisons using Bonferroni correction (3 trial epochs x 2 trial types). Our rationale for measuring Ca^2+^ spike activity as a point process was that events were often loosely time-locked with the onset or offset of specific trial epochs, which complicates any interpretation of trial-averaged fluorescence signals. For analyses in Figures 2-4 and 9-10, mice’s behavioral performance was >70% (mean d-prime = 2.72 ± 0.08) and only correctly answered trials were analyzed so as to determine if dendritic activity varies with trial type, rather than trial outcome (Mean trial counts: 350.0 ± 9.6. Range: 169-657. n = 78 sessions). Dendrites were classified as sound excited or inhibited if the Ca^2+^ spike probability during sound presentation was significantly greater or lower than during the 2-3 s pre-sound baseline period, respectively. Dendrites were classified as early answer responsive if they displayed an increase in Ca^2+^ spike probability relative to baseline 1 s following sound offset. Late answer responsive dendrites were defined as ROIs which showed a significant increase in Ca^2+^ spike probability relative to baseline late in the trial period, 2-4 s following sound offset.

In Figure 3, the Selectivity Index for each dendrite was calculated as

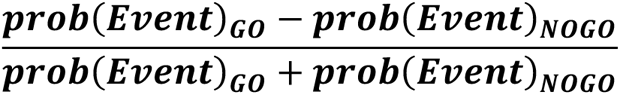

Analyses in Figures 5-7 were run on a subset of sessions where mice performed above chance but made enough false alarms on NO-GO trials for meaningful comparisons (mean false alarm rate: 26.8 ±1.5%, range 15.8 – 51.7%, n = 38 sessions). We did not include activity on miss trials in these analyses as mice typically had few, if any misses on GO trials during task engagement.

In Figures 12-13, behavioral trials were discarded for analyses after mice made 5 or more consecutive misses on GO trials, as mice were assumed to be satiated and no longer motivated to perform the task. Both groups of mice performed a similar number of trials until satiety (mean: 284.4 ± 2.9 vs 277.1 ± 3.6; median: 300 vs 300 trials for control and virus groups, respectively. p=0.56, Kolmogorov-Smirnov test).

Although not explicitly determined *a priori*, the sample sizes employed in this work are consistent with established standards in the field. For all within- and between-group comparisons, data were tested for normality using Lilliefors test prior to significance testing. Normally distributed data were compared via parametric tests, whereas non-parametric approaches were employed when one or more of the datasets deviated from normality. Mean ± SEM and median are reported for normally and non-normally distributed data, respectively. Alpha value (<0.05) is corrected for multiple comparisons when appropriate.

### Histology and confocal imaging

Mice were euthanized via an overdose of isoflurane within an induction chamber. Transcardial perfusions were performed after breathing and reflexes were absent using PBS followed by 10% buffered formalin. Brains were carefully removed and stored in the dark in 10% formalin overnight at 4°C for immersion fixation. Prior to vibratome slicing, brains were rinsed 3 times over 5 minutes in PBS on a platform shaker and embedded into 4% agar (PBS). Coronal slices (100 µm) were cut using a Leica vibratome (VT1000S) and chilled PBS. Slices containing the IC and auditory cortices were mounted on object slides and covered using Fluoromount-G (SouthernBiotech). Slides were dried at room temperature overnight, sealed using clear nail polish and stored in the dark at 4°C until confocal imaging. Images were collected using a Leica TCS SP8 laser scanning confocal microscope equipped with a 10x objective.

## Results

### Head-fixed mice discriminate sound temporal envelope fluctuations in a GO/NO-GO paradigm

Water-deprived, head-fixed mice (n=14) were trained to report the presence or absence of sinusoidal amplitude-modulation (**sAM**; 15 Hz modulation rate, 100% modulation depth, 1 s duration) in a 2-octave band-limited noise carrier (4-16 kHz; 65-70 dB SPL). On 50% of trials, a GO sound instructed that licking a waterspout during an answer period following sound offset would deliver a drop of 10% sugar water. In the other half of trials, a NO-GO sound instructed to withhold licking; incorrect licks during the answer period (false alarms) were punished with an increased inter-trial interval (Figure 1A). Whether the sAM sound served as GO or NO-GO was counterbalanced across mice (n=7 mice in each group). Mice rapidly learned associations between the GO sound and reward delivery, as evidenced by a consistently high hit rate even during the first five training sessions (Figure 1B). The number of false alarms gradually fell across training, such that mice’s sensitivity index (d-prime; Tanner and Swets, 1954) increased with continued training (Figure 1B,C); this effect was primarily due to an increased propensity of withholding responses on NO-GO trials (Figure 1D,E). Task acquisition was also associated with a reduced licking during the pre-sound baseline period in expert mice, and lick bout initiation became increasingly limited towards the end of the GO sound (Figure 1F). Consequently, subjects’ “accuracy” of lick bouts limited to the answer period increased across training (Figure 1G), similar to previous studies in head-fixed mice (Komiyama et al., 2010). Altogether these data show that head-fixed mice rapidly learned to attend to, and subsequently discriminate, the temporal envelope of sounds.

**Figure 1:**
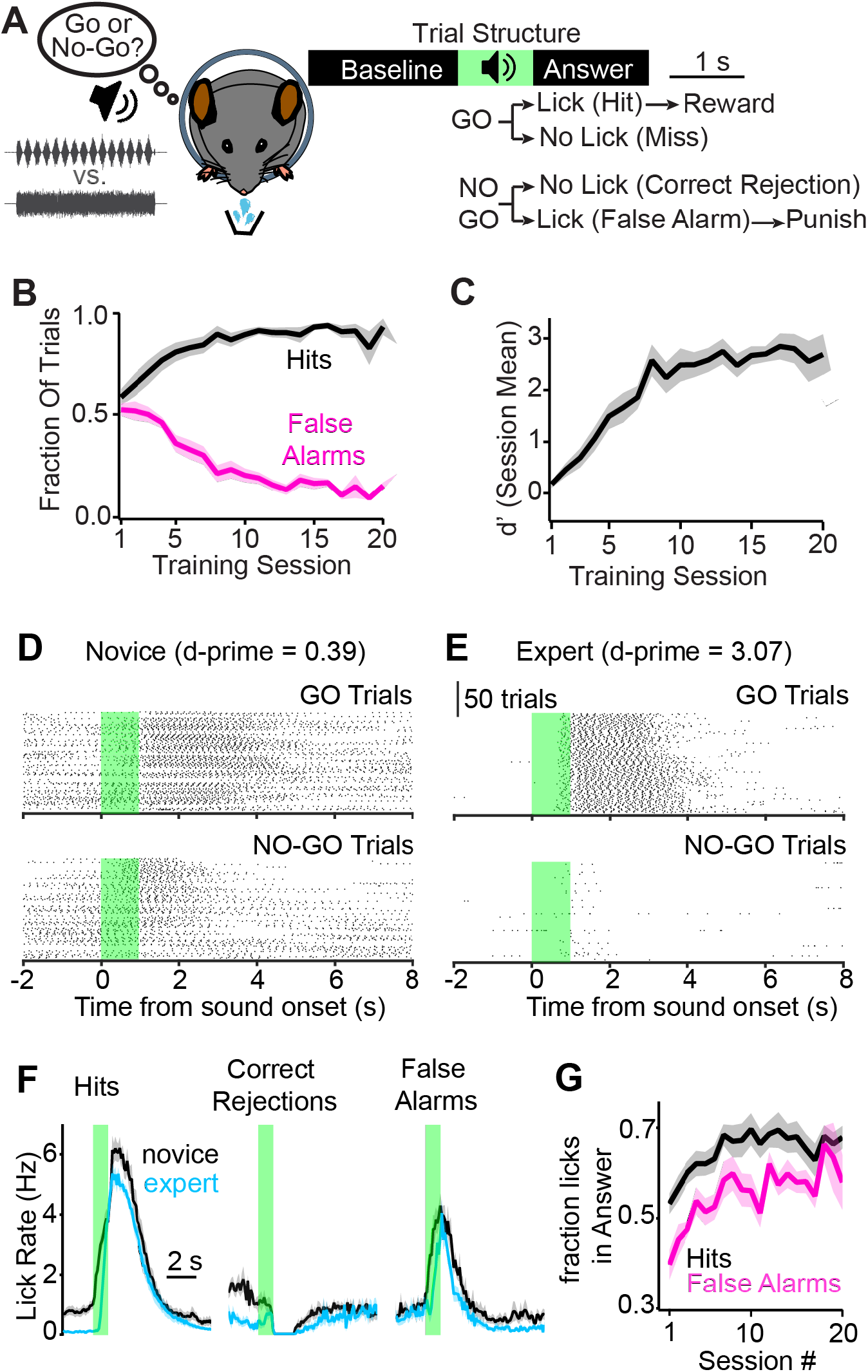
Head-fixed mice discriminate temporal envelope fluctuations in a GO/NO-GO paradigm. **A)** Cartoon and trial structure. **B)** Mean hit (black) and false alarm (magenta) rates across learning for n = 14 mice. **C)** Similar to panel B, but for the sensitivity index (d-prime). **D,E)** Example lick rasters from a single mouse during its first (novice; D) and last (expert; E) training session. Green bars denote sound presentation. Of note is the high rate of licking during the baseline period before sound onset during the novice, but not expert session. Data in D and E are from the same mouse. **F)** Lick PSTHs in novice and expert sessions, averaged across n = 14 mice and grouped by trial outcome. **G)** Summary data plotting the mean fraction of licks in the answer period on Hits (black) and False Alarms. A mixed effects model revealed a main effect of session # (F(4.74,61.61)=4.602, p=0.0015), trial outcome (F(1.0,13.0)=25.78, p=0.0002), but no a session # x trial outcome interaction (F(3.40,24.40)=0.604, p=0.19).

### Diverse activity profiles of auditory corticofugal dendritic spikes in behaving mice

The IC of the midbrain receives major descending projections from the auditory cortex, with 80-90% of the auditory cortico-collicular pathway arising from thick-tufted layer 5 pyramidal neurons (Winer et al., 1998; Schofield, 2009; Slater et al., 2013; Williamson and Polley, 2019; Yudintsev et al., 2021; Lesicko et al., 2022). Thus, we expressed the Ca^2+^ indicator GCaMP6f and a nuclear localized tdTomato in auditory corticofugal neurons via a retrograde AAV injected into the left IC (Figure 2A). Labeling in auditory cortex was restricted to layer 5 pyramidal neurons (Figure 2B) and we did not observe any labeling in the 10-20% of layer 6 auditory cortico-collicular neurons. This layer 5 specific tropism is expected given the known properties of rAAV2-retro (Tervo et al., 2016).

**Figure 2:**
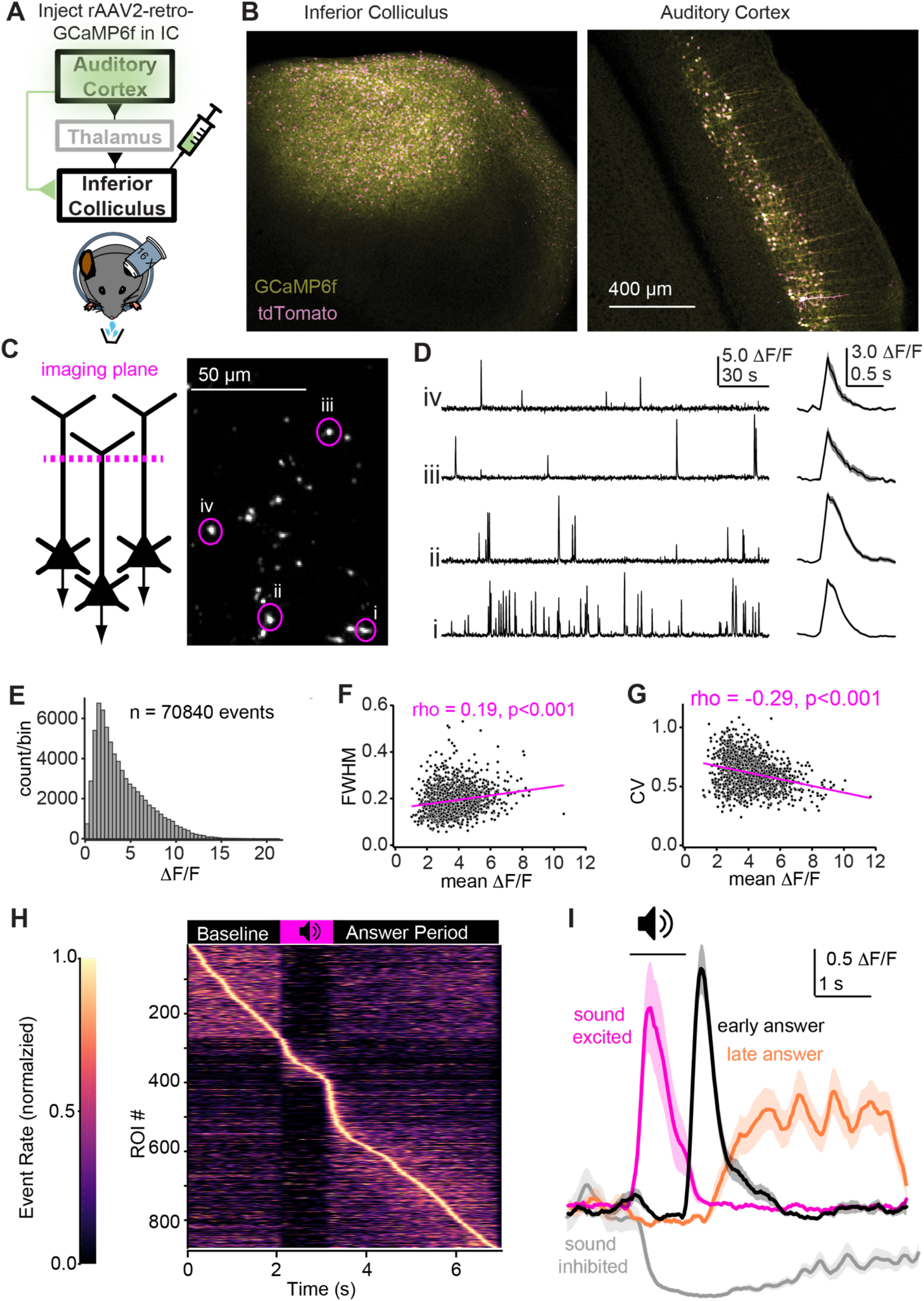
Dendritic Ca^2+^ spikes in auditory corticofugal neurons of behaving mice. **A)** Cartoon of experiment. A retrograde virus bearing the GCaMP6f payload is injected in the IC; imaging data are collected from the auditory cortex of head-fixed mice during performance of the task in Figure 1. **B)** Confocal micrographs of the IC (Left) and auditory cortex (Right) show preferential retrograde labeling of layer 5 pyramidal neurons. **C)** Trunk dendrites are imaged ∼150-300 µm below the auditory cortical surface. Micrograph (right) is an example FOV with dendrite cross-sections visible as circular or ovoid structures; i-iv denote example ROIs. **D)** Fluorescence time series from ROIs in panel F. Dendritic spikes are observable as large and fast transients. Right: Mean Ca^2+^ spikes in each ROI after onset alignment of individual transients. E) Distribution of peak amplitudes for 70,840 dendritic spike events. F,G) correlation between the mean peak ΔF/F (x-axes) and either half-width (F) or coefficient of variation (G) across all dendrites. **H)** Normalized event rates of n = 880 task modulated dendrites. Of note is the paucity of dendrites with peak rates during sound presentation. **I)** Example averages for distinct type task-modulated dendrites. Shading in all panels is ±SEM.

In layer 5 pyramidal neurons, Ca^2+^ fluorescence signals in distal apical trunk dendrites specifically reflect locally generated Ca^2+^ spikes rather than Na^+^ action potentials back-propagating from the axosomatic compartment (Pérez-Garci et al., 2006; Xu et al., 2012a; Harnett et al., 2013; Beaulieu-Laroche et al., 2019). Thus, we measured dendritic spikes in auditory corticofugal neurons using 2-photon imaging of apical trunk dendrites 150-300 µm below the pial surface (Figure 2C). We quantified the activity of 1554 apical trunks in 78 sessions from 14 mice engaged in the GO/NO-GO task. GCaMP6f signals in dendritic regions of interests (ROIs) were of large amplitude and displayed fast kinetics (Figure 2D,E. Median half-width and amplitude: 183 ms and 3.7 ΔF/F). Moreover, there was a weak but significant correlations between dendritic spike amplitude and half-width (Figure 2F), as well as the coefficient of variation in amplitude (Figure 2G). These results are similar to previous reports of GCaMP6f signals from layer 5 trunk dendrites in somatosensory cortex (Ranganathan et al., 2018), implying that auditory corticofugal GCaMP6f signals are mediated by similar mechanisms.

56.7% (n=880) of dendrites showed task-related modulation during task performance, defined as a statistically significant change in Ca^2+^ event rate during the sound or answer period on correctly answered GO and/or NO-GO trials. Interestingly, only ∼11.9% (n = 105) of task-modulated dendrites showed activity increases during the sound cue (Figure 2H; Figure 2I, magenta). Instead, ∼48.3% (n=425) of dendrites showed a *reduction* in Ca^2+^ events during the sound, thereby reducing the mean fluorescence relative to baseline (Figure 2I, grey). These data suggest that in a large fraction of corticofugal neurons, the behaviorally relevant sounds used in our paradigm powerfully reduce dendritic electrogenesis. In addition, ∼45% of dendrites (n=398) were active in the answer period either during the first second following sound offset (early answer responses; n = 98; Figure 2I, black), for several seconds during the inter-trial interval (late answer responses; n = 185; Figure 2I, orange), or during both early and late answer epochs (n = 115). During active listening, dendritic spikes in auditory corticofugal neurons may be anti-correlated with behaviorally relevant sounds. Moreover, only ∼34% of sound excited (n = 36/105) and ∼18% of sound inhibited dendrites (n = 78/425) also increased their activity during the answer period, implying that the different activity patterns of corticofugal neurons in our task reflect largely non-overlapping populations.

### Sound and answer responsive dendrites are preferentially active on GO trials

The distinct activity profiles of corticofugal neurons may reflect preferential responses on GO or NO-GO trials (O’Connor et al., 2010). We tested if activity was differentially modulated during the sound and answer period of different trial types by calculating a selectivity index for each task-modulated dendrite. Values of +1 and -1 indicate that dendritic spiking is exclusively modulated on GO and NO-GO trials, whereas a value of 0 indicates identical modulation on each trial-type. The population distribution of selectivity indices in sound excited dendrites was positively skewed, e.g. showed a strong bias towards GO trials (Figure 3A; median selectivity index = 0.56). This result was not due to an innate preference of dendritic spikes for particular acoustic features, as the distribution of selectivity indices was similar across mice trained with the sAM sound as the GO and NO-GO stimuli (p=0.998, Kolmogorov-Smirnov test).

**Figure 3:**
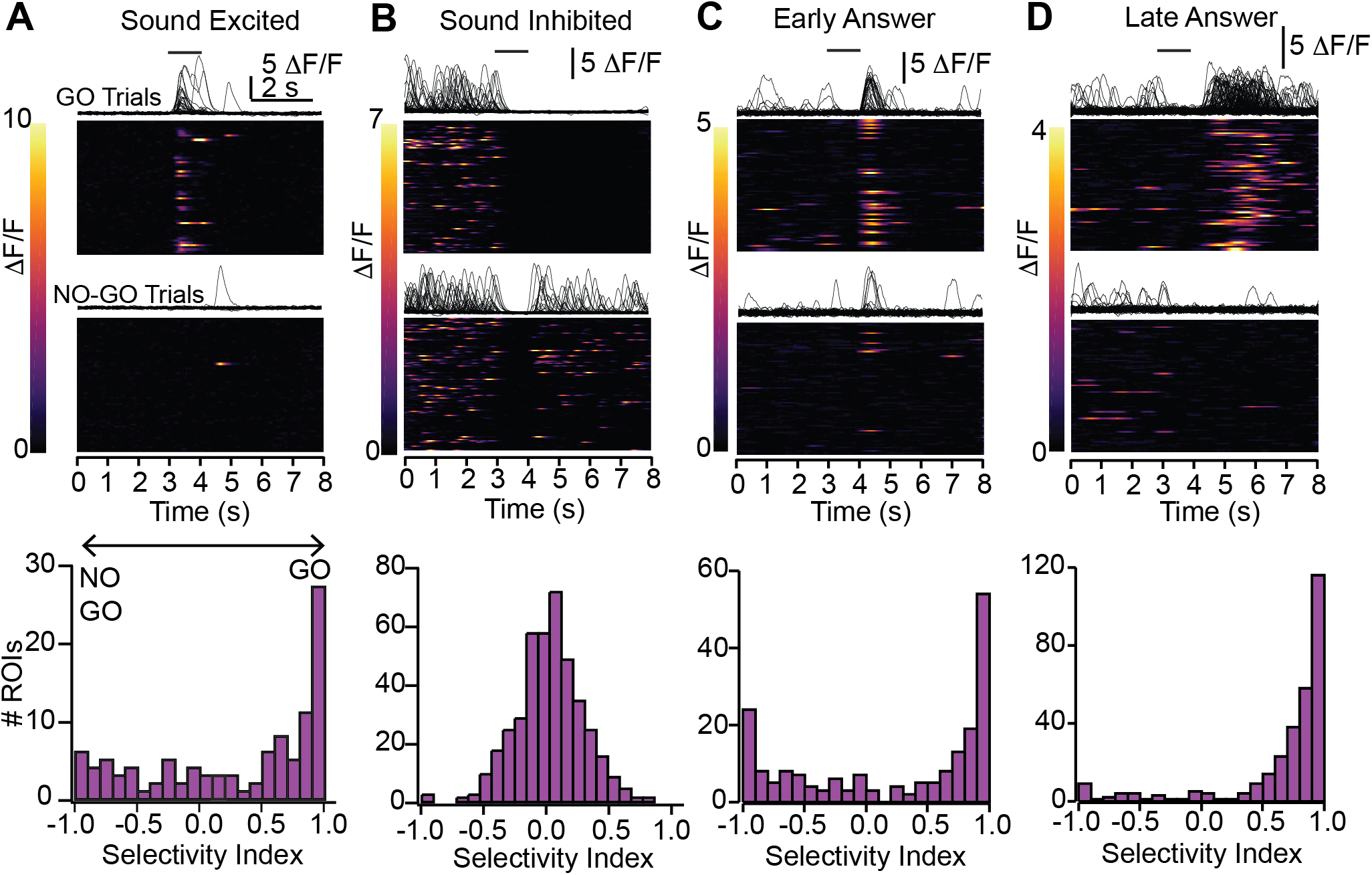
Dendritic spikes are biased towards GO trials. **A)** Example heatmaps from correctly answered GO and NO-GO trials in a sound-excited dendrite are shown in the upper panels, with corresponding raw traces. Black bar denotes sound onset The lower panel histogram shows the distribution of selectivity indices (see Results and Methods) across the population of sound excited dendrites. B-D are organized same as panel A, but for sound inhibited (**B**), early answer (**C**) and late answer dendrites (**D**).

Conversely, selectivity indices of sound inhibited dendrites centered near 0 (median = 0.0253, Figure 3B), reflecting a powerful and non-specific reduction of dendritic spikes during the discriminative sounds. In addition, 40% of sound inhibited dendrites (n=172) also displayed sustained inhibition during the answer period on GO trials as well as on false alarm trials (Figure 3B), suggesting that the mechanisms responsible for reducing dendritic excitability are engaged during both sound presentation and the answer period.

Both early and late answer responsive dendrites had positively skewed selectivity indices similar to sound excited dendrites (Figures 3C,D. Median values of 0.61 and 0.85, respectively), indicating that answer period activity occurs primarily (though not exclusively) on GO trials. These data show that dendritic spikes preferentially occur on trials involving reward predictive sounds and reward-related licking. One possibility is that these answer period responses reflect motor- or reward-related activity, or alternatively, arousal-related neuromodulatory transmission which triggers dendritic spikes *in tandem* with reward delivery and/or mice’s actions (Labarrera et al., 2018; Williams and Fletcher, 2019). Alternatively, the GO selectivity could also reflect learned acoustic responses biased to the termination of reward predictive sounds (Lee and Rothschild, 2021).

### Corticofugal dendrites with similar task responsive activity are not spatially clustered

Sensory cortices often exhibit topography: Neurons with similar receptive fields are spatially clustered within and/or across cortical layers (Hubel and Wiesel, 1963; Merzenich et al., 1975; Issa et al., 2014; Peron et al., 2015; Schmitt et al., 2023 but see Bandyopadhyay et al., 2010; Rothschild et al., 2010). We therefore asked whether corticofugal neuron also formed spatial clusters depending on whether they exhibited similar or distinct task-relevant activity patterns. If this hypothesis is correct, pairwise Euclidean distances in a given FOV should be shorter when measured between dendrites within the same task-response category compared to values measured across task-response profiles. However, pairwise Euclidean distances between sound-excited dendrites were not significantly different than between sound excited and sound inhibited dendrites, or between sound-excited and early/late answer responsive dendrites (Figure 4A; Kruskal-Wallis test, KW statistic: 6.56, p=0.16). Similarly, the distribution of pairwise Euclidean distances for sound-inhibited (Figure 4B; Kruskal-Wallis test, KW statistic: 7.13, p=0.13), early answer (Figure 4C; Kruskal-Wallis test, KW statistic: 7.365, p=0.118), or late answer responsive dendrites (Figure 4D; Kruskal-Wallis test, KW statistic: 5.59, p=0.13) were not significantly different when tested within and across task-response categories. Altogether these data suggest that task-relevant dendritic activity does not display overt topography, at least on the spatial scale of our FOVs (<500 μm^2^).

**Figure 4:**
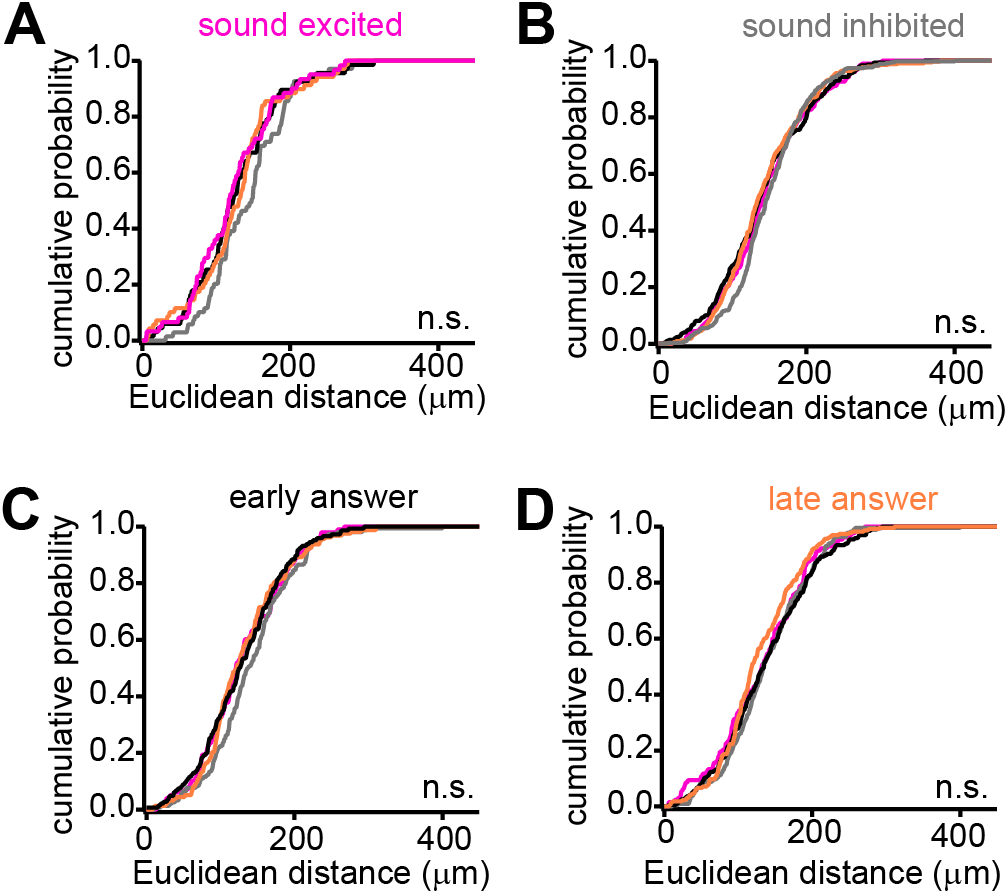
ROIs with different response profiles do not appear spatially clustered. **A)** Normalized cumulative probability distributions of pairwise Euclidean distances between sound excited dendrites and sound inhibited, early answer, late answer, or other sound excited dendrites (gray, black, orange, and magenta, respectively). **B-D)** Same as panel A but for sound inhibited (B), early answer (C) or late answer (D) responsive dendrites. Of note are the similar distributions of data across all comparisons, indicating that task-responsive dendrites do not exhibit noticeable spatial clustering.

### Answer selectivity on GO trials occurs in tandem with goal-directed licking, independent of reward consumption

The biased selectivity of dendritic spikes during the answer period of GO trials could encode the termination of a behaviorally relevant sound features. Indeed, the temporal precision of early answer responses (Figure 3C) is reminiscent of auditory cortical “OFF” responses (Liu et al., 2019; Solyga and Barkat, 2021) that are known to be patterned by the learned salience of sounds (Lee and Rothschild, 2021). If this hypothesis is correct, GO selective dendrites should have distinct activity profiles on GO and NO-GO trials where different sound cues are presented, regardless of mice’s behavioral choice during the answer period (Figure 5A, left).

**Figure 5:**
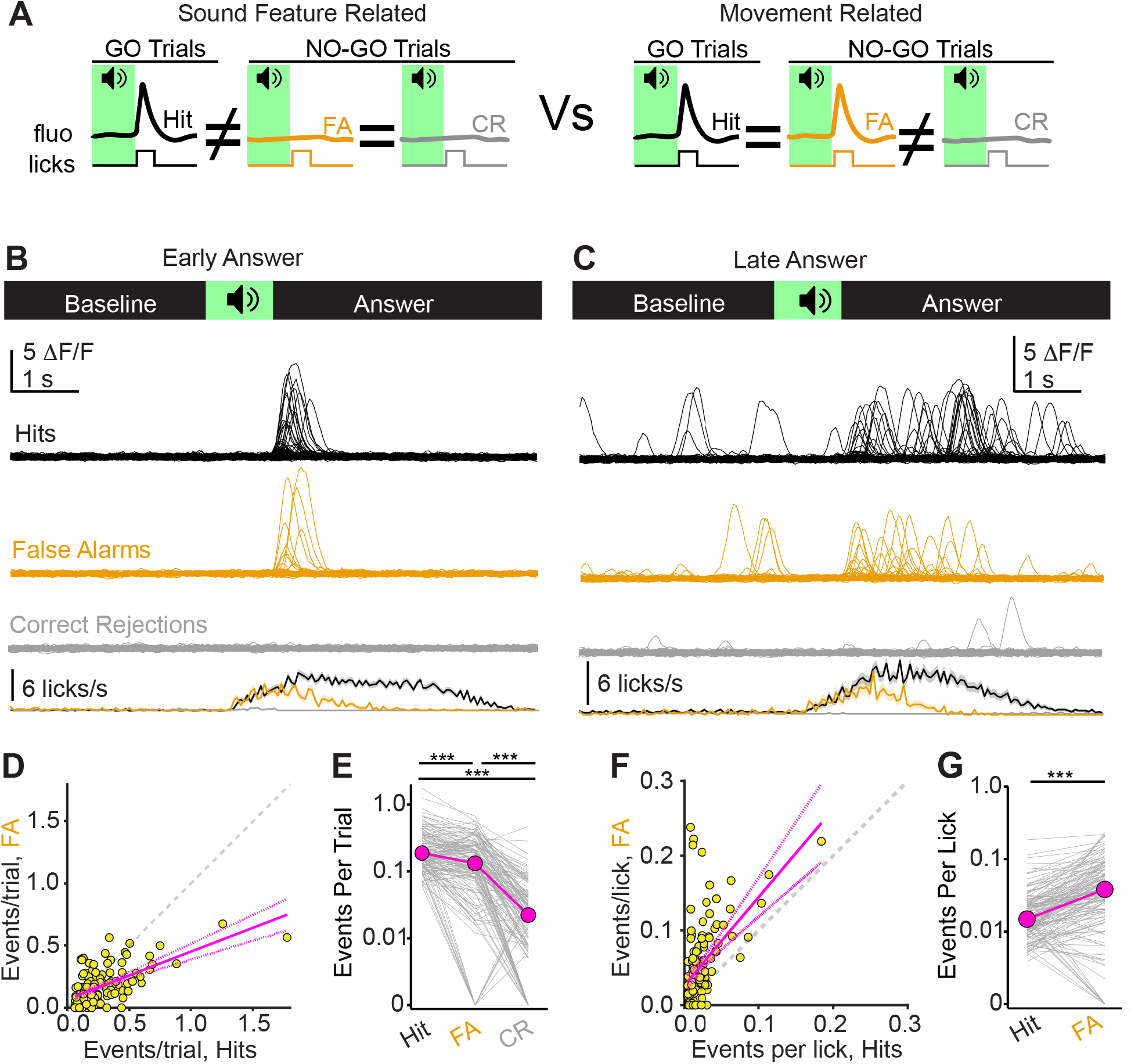
GO-selectivity in answer responsive dendrites correlates with goal-directed actions. **A)** Cartoon of analytical rationale. **B)** Example traces and lick rate histograms (upper and lower panels, respectively) from an early answer responsive dendrite. Of note are dendritic spikes occurring on hits and false alarms, but not on correct rejections. The minor number of lick events in the histogram of Correct Rejections reflects occasional early lick bouts that the mouse subsequently terminated prior to the answer period (see also Methods, *Behavior during 2-photon Ca^2+^ Imaging)*. **C)** Same as panel B, but for a late answer responsive dendrite. **D)** The number of dendritic spikes per trial on false alarms versus hits (y and x axes, respectively) for answer responsive dendrites. Magenta is the linear fit with 95% confidence intervals. The gray dotted line is the unity. **E)** Events per trial across the three trial outcome conditions for answer responsive dendrites. Gray lines are individual ROIs, magenta is median. **F)** Same as D, but normalized to the number of licks on each trial. Most points fall above the unity line, indicating more dendritic spikes per lick on false alarms compared to hits. **G)** Events per lick on hits and false alarms. Asterisks denote statistical significance. Of note, y-axes for E and G are on a log scale. Early and late dendrites are pooled in all analyses. *** = p<0.0001

Alternatively, answer period activity could reflect movement-related activity during reward consumption (Huang et al., 2019; Lacefield et al., 2019), arousal-related increases in neuromodulatory transmission during operant actions (Labarrera et al., 2018), or perhaps exertion of physical force (Bakhurin et al., 2023). In these scenarios, answer period activity should be similar on hit and false alarm trials; mice would hear different sounds but perform similar operant actions (Figure 5A, right). Most GO selective, answer responsive dendrites were active on both hit and false alarm trials and showed far less activity on correct rejection trials; this result held true across early and late answer populations (Figure 5B,C). Additionally, there was a significant positive correlation between dendritic spike probability on hit and false alarm trials (Figure 5D; Pearson’s rho =0.62, p<0.001). Thus, dendritic spikes during the answer period of GO trials co-occur with mice’s instrumental actions, and are not explicitly triggered by sound termination.

Indeed, dendritic spike probability was generally higher on hit compared to false alarm trials (Figure 4E. Median event probability per trial: 18.8, 13.2 and 2.2% for hits, false alarms, and correct rejections, respectively. F(3,137)=176.8, p<0.0001, Friedman test. P<0.0001 for all pair-wise comparisons, Dunn’s post-hoc tests). However, this result likely reflects differences in the absolute number of licks mice made on each trial type: Normalizing the dendritic spike probability to mice’s lick rate revealed a *greater* probability of dendritic spikes on false alarm compared to hit trials (Figure 5F,G. Median events per lick: 0.015 vs. 0.038 for hit and false alarm trials, respectively. Signed-Rank statistic = 924, p<0.0001, signed-rank test). The data suggest either a ceiling effect for dendritic spikes that occur during licking, or alternatively, that reward consumption has a net inhibitory effect on dendritic electrogenesis. Furthermore, the peak fluorescence of individual dendritic events was similar on hits and false alarm trials, implying that the probability, but not the magnitude of dendritic spikes co-varies with mice’s actions (Median peak ΔF/F of 3.37 vs. 3.47 for hits and false alarms, respectively. p=0.58, signed-rank test). *In tandem* with the results of Figure 3, our data imply that the selectivity of dendritic spikes towards GO trials reflects two mechanisms: In sound excited dendrites, a learning-dependent process biases dendritic responses towards behaviorally relevant sound features, whereas answer period activity is mostly correlated with the initiation and continued execution of goal-directed movements, a process which may or may not be patterned by learning.

Although answer period activity was largely skewed towards GO trials, a minor population of dendrites showed heightened activity specifically in the answer period of correctly answered NO-GO trials. Activity of these dendrites thus correlated with mice’s choice, but occurred preferentially on correct rejection rather than false alarm trials (Figure 6A-D. See also Figure 3C,D; n = 26 dendrites). Consequently the median event probability was 5.0, 12.2, and 18.5% for hits, false alarms, and correct rejections, respectively. (Figure 6E. F(3,31)=28.31, p<0.0001, Friedman test. P<0.01 comparing correct rejections vs hits or false alarms, p=0.0925 hits vs false alarms, Dunn’s post-hoc tests). One potential interpretation is that this NO-GO selectivity reflects high-level signals related to the withholding of licking. Alternatively, selectivity for correct rejections may reflect dendrites that are broadly tuned to sound termination, and the reduced activity on false alarm trials could arise from inhibitory inputs that whose activity correlates with mice’s licking action during the answer period.

**Figure 6:**
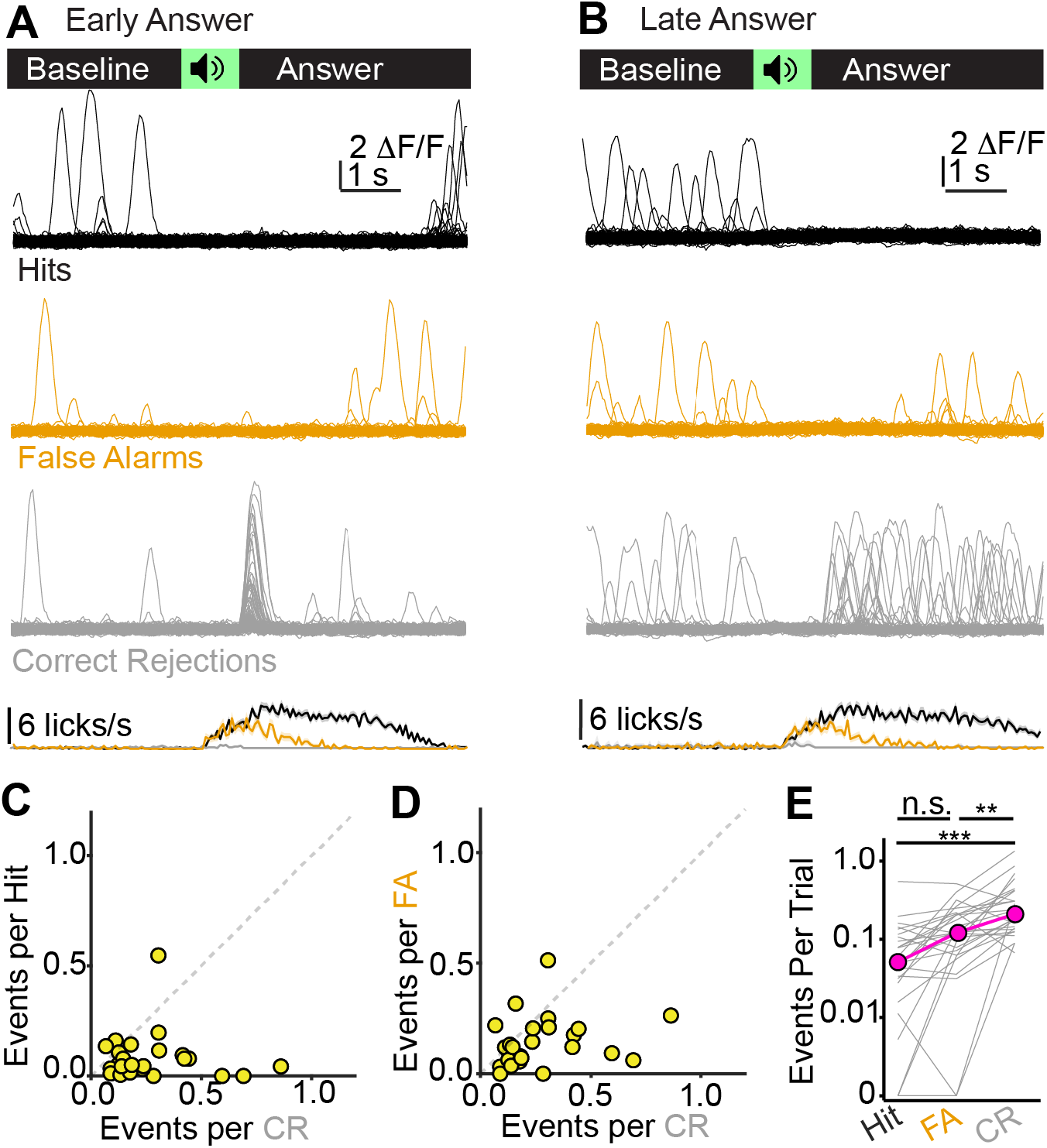
NO-GO selective, answer responsive dendrites are preferentially active on correct rejection trials. **A)** Overlaid fluorescence traces and lick rate histogram for a NO-GO selective, early answer responsive dendrite on hits, false alarms, and correct rejections. This ROI was recorded in the same session as the example early answer dendrite in 4B, such that the behavioral traces are identical. **B)** Same as panel A, but for a NO-GO selective late answer responsive dendrite. Of note in A and B is that the increase in dendritic Ca^2+^ spikes during the answer period does not occur on false alarm trials. **C)** Number of dendritic spikes on hits and correct rejections (y and x axes, respectively) are plotted for n = 26 NO-GO selective, answer responsive dendrite. **D)** Same as C, but for false alarms and correct rejections. Of note is that most datapoints fall below the unity line in both panels, indicating selective activity on correct rejections. **E)** Summary data showing the number of events per trial for each outcome category. Of note, y-axis is a log scale. ** = p < 0.01, *** = p<0.0001.

### Sound excited dendritic activity does not co-vary with trial outcome

We next asked if the activity of sound-excited dendrites similarly co-varied depending on trial outcome. Indeed, activity of GO preferring, sound-excited dendrites may not explicitly reflect tuning to specific acoustic features, but rather could correlate with mice’s instrumental actions as in Figure 5. Similarly, the subset of NO-GO preferring, sound-excited dendrites might correlate with mice’s withholding of licking as in Figure 6. Activity of sound-excited dendrites would correlate with trial outcome, rather than trial type, and as such should be more similar on hit and false alarm compared to hit and correct rejection trials; this prediction should hold true regardless of whether a given dendrite is preferentially active during sound presentation of GO or NO-GO trials. Alternatively, a result showing that activity is similar on divergent trial outcomes of the same trial type would argue that dendritic responses during sound presentation reflect acoustic features. In agreement with the latter prediction, we found that activity of GO preferring dendrites occurred preferentially on hit trials, with similarly sparse activity on correct rejection and false alarm trials (Figure 7A). NO-GO preferring dendrites showed similar activity pattern on correct rejections and false alarms, with minimal activity on hit trials (Figure 7B). Peri-stimulus time histograms of dendritic activity on different trial outcomes revealed greater activity on hit compared to correct rejection and false alarm trials, in agreement with our observation that activity of sound-excited dendrites is biased towards GO trials (Figure 7C). To explicitly test if dendritic activity is more closely related to trial outcome than sound features, we calculated the absolute activity difference across trial timepoints between hit and correct rejection, as well as hit and false alarm trials for all sound-responsive dendrites in these datasets. The two curves were similar (Figure 7D), and a two-way, repeated measures ANOVA revealed a main effect of timepoint (F(27.3,1338)=10.22, p<0.0001), but no main effect of trial-type comparison (F(1.0,49.0)=2.80, p=0.1) or trial-type comparison x time-point interaction (F(31.46,1542)=1.137, p=0.276). Thus, absolute activity differences are similar between hit trials and NO-GO trials with divergent outcomes, thereby arguing that activity in sound-excited dendrites correlates with the identity of the discriminative sound cue rather than mice’s actions.

**Figure 7:**
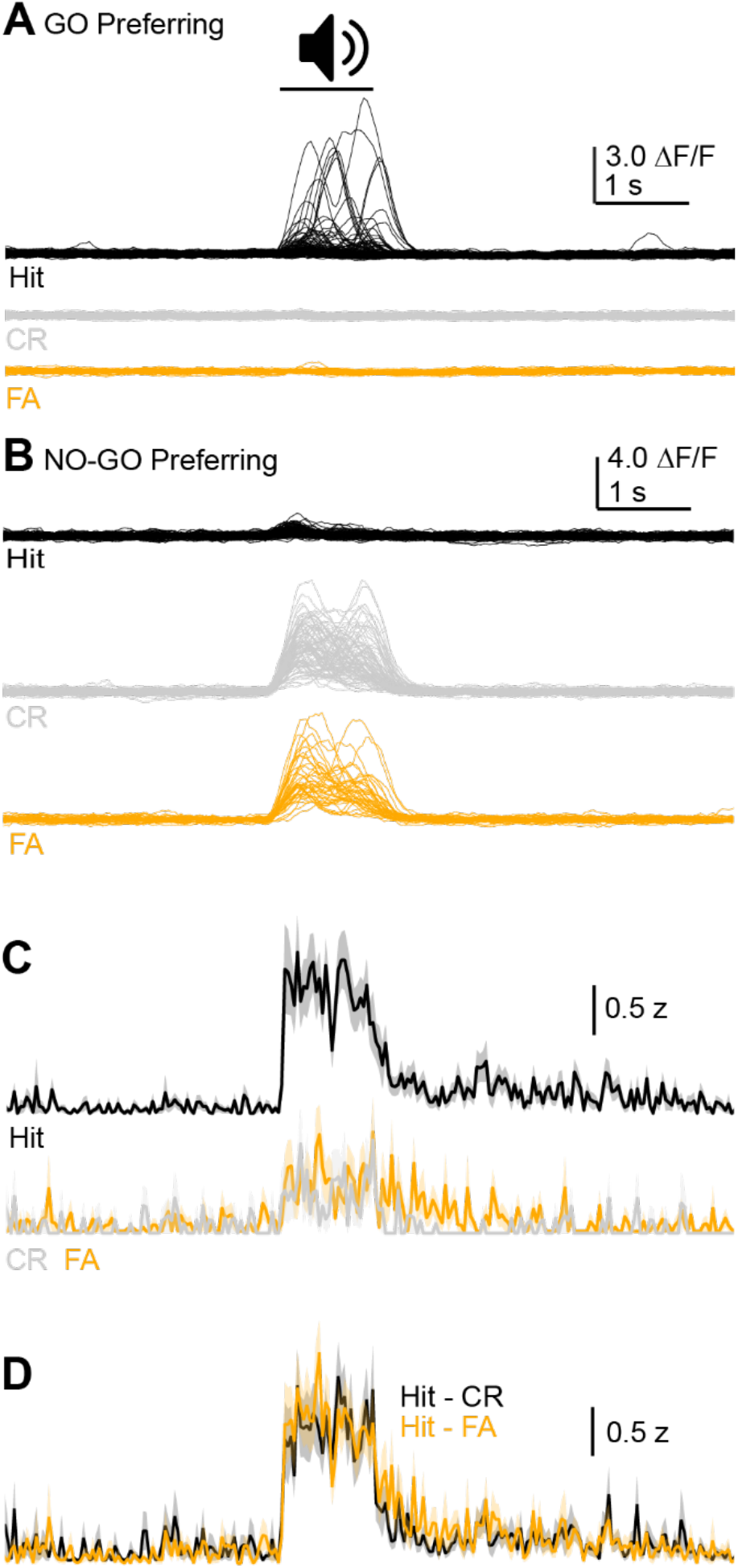
Sound excited dendrite activity does not co-vary with trial outcome. **A)** Example single trials for a GO-preferring, sound-excited dendrite. Of note is the strong activity on Hit trials (black traces), and similar lack of responses on Correct Rejection and False Alarm trials (gray and orange, respectively). **B)** Same as panel A but for a NO-GO preferring dendrite. **C)** Mean PSTH for dendritic events on Hit, Correct Rejection, and False Alarm trials for n = 50 sound excited dendrites. Data from GO and NO-GO preferring dendrites are pooled. Color scheme is same as in panels A and B. Shading is ± SEM. **D)** The difference in activity between hit and correct rejection (black) or hit and false alarm (orange) trials is plotted across time for the dendrites in panel C. Of note is that the curves are similar in the two conditions, indicating that neural activity in sound-excited dendrites co-varies with the sound stimulus rather than trial outcome.

### Licking-related dendritic activity does not require task learning or engagement

Previous data suggest that the motor- and/or reward-related activity in sensory cortices may arise as a consequence of learning (Lacefield et al., 2019; Takamiya et al., 2022). By contrast, non-sensory and putatively motor-related activity also occurs in non-task engaged animals (Schneider et al., 2014; Stringer et al., 2019). Does the dendritic activity we observe in the answer period require task learning and/or engagement, or can similar activity patterns be evoked in absence of learning? We tested this idea in a separate cohort by imaging auditory corticofugal dendrites while untrained, water-deprived mice consumed sugar water droplets delivered passively through a reward spout (Figure 8A). Interestingly, a subset of corticofugal dendrites showed clear increases in dendritic spike probability when mice licked the water spout following reward delivery (Figure 8B; n = 29/232 dendrites in N = 5 mice), with the majority (79 +/- 2%) of dendritic events in these sessions occurred within 4 s of reward delivery. Thus although learning may increase the prevalence of non-auditory signals in primary sensory cortical fields, substantial activity co-occurs with mice’s actions even under non-task engaged conditions and thus does not necessarily reflect learned information.

**Figure 8:**
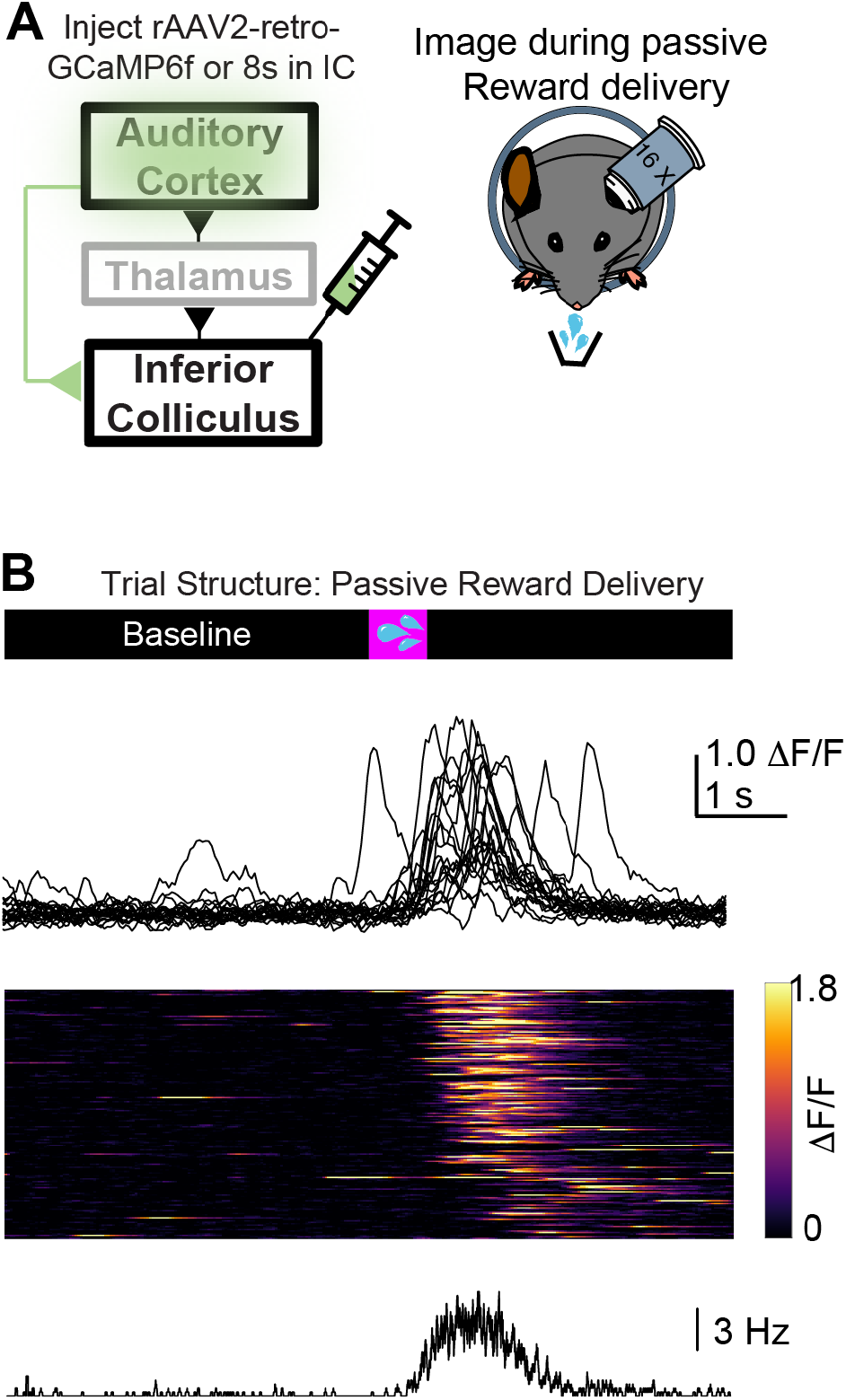
Mice’s licking during passive reward delivery triggers dendritic Ca^2+^ events. **A)** Experimental design: rAAV2-retro virus packaging GCaMP6f or 8s is injected in the IC (N = 3 and N = 2 mice expressing 6f and 8s, respectively). Imaging is performed in auditory cortex during passive reward delivery. **B)** Example dendritic ROI showing responses to mice’s licking during passive reward delivery. Upper panel shows overlaid single trial fluorescence (data are from a GCaMP8s expressing dendrite). Middle panel is heatmap of all trials from this ROI in a single session; lower panel is averaged lick PSTH during this session. Similar results were observed in n=29/232 ROIs from n=5 mice.

### The onset of answer period activity is yoked to the first lick following sound termination

Mice’s lick bouts often began towards the end of the discriminative sound (Figure 1D-F; 5B,C; 6A), whereas the increased activity of answer responsive dendrites often lagged this anticipatory licking and appeared restricted to trial periods following sound termination (Figure 3C,D; 5B,C). This observation suggests that acoustic inputs may have a net inhibitory effect on licking-correlated dendritic spikes.

To test this hypothesis, we aligned the fluorescence data of GO selective, answer responsive dendrites to the first lick on each trial occurring either during sound presentation, or the answer period (n = 317 dendrites; Figure 9A and 9B, respectively). Trial-by-trial inspection of the raw data showed that dendritic spikes occurred with variable latencies following the first lick during sound presentation, and that the shortest latencies were on trials where mice’s first lick occurred towards the end of the sound (Figure 9A; compare yellow and purple traces). By contrast, dendritic spike events were time-locked to the first lick in the answer period following sound offset, regardless of whether the first lick occurred early or late in the answer period (Figure 9B). A qualitatively similar phenomenon can be observed in group-averaged fluorescence traces: The fluorescence increase occurs *in tandem* with the first lick during the answer period (Figure 9C, magenta), whereas it lags the first lick during sound presentation by ∼500 ms (Figure 9C, black).

**Figure 9:**
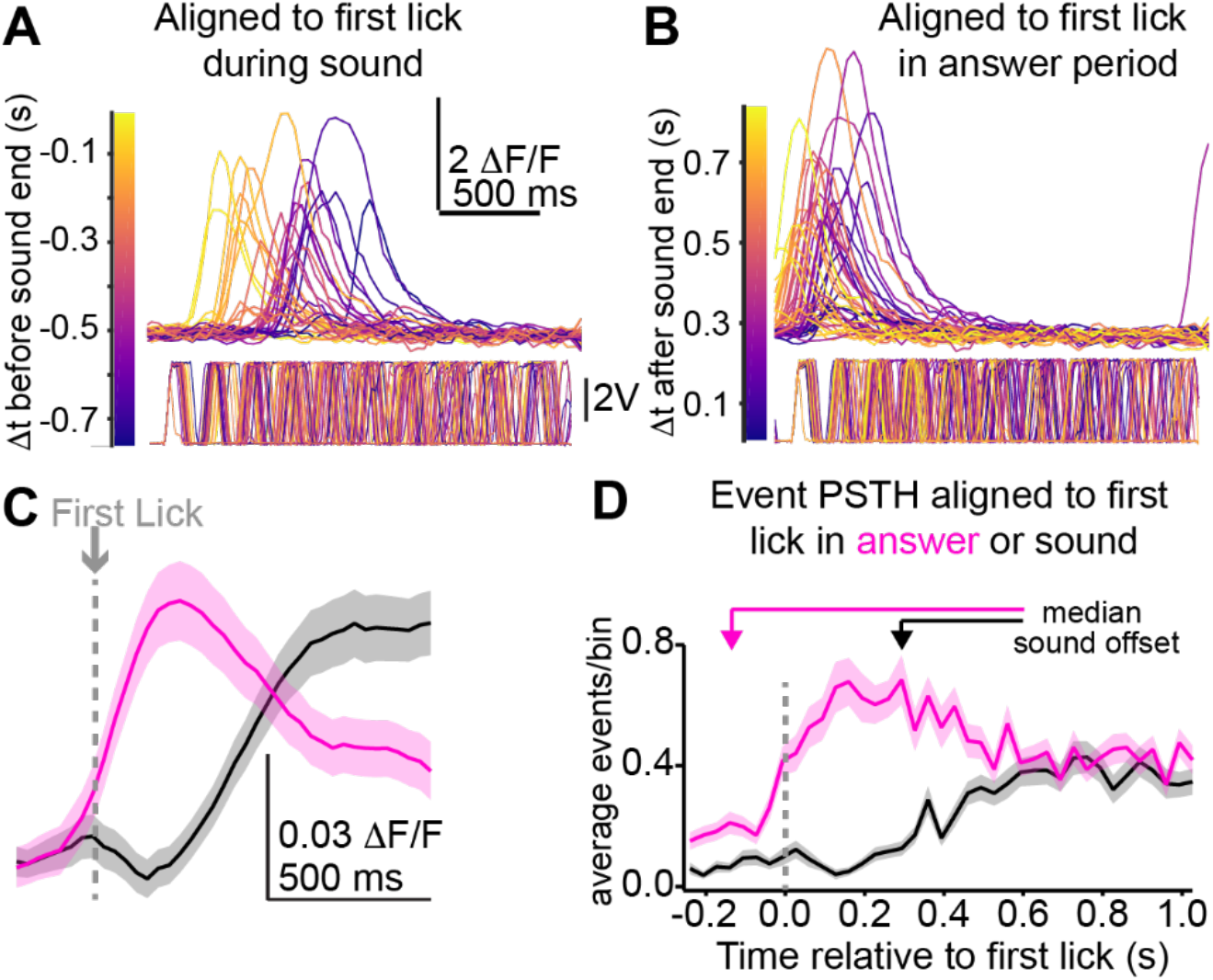
Answer period activity is preferentially triggered by licks occurring following sound offset. **A)** Example fluorescence traces from a single dendritic ROI, aligned to the first lick occurring during sound presentation. The traces are color coded by the latency from first lick event to sound termination. Lower panel is lick sensor voltage recording. Of note, the latency between the first lick and onset of Ca^2+^ spike varies on a trial-to-trial basis, with shorter latencies observed when first licks occur closer to sound termination (compare yellow and purple traces). **B)** Same as panel A, but with fluorescence traces aligned to mice’s first lick in the answer period following sound offset. Color coding reflects the time difference between sound termination and first lick event. Of note is the lower onset variance in Ca^2+^ spike onset compared to panel A. **C)** Summary fluorescence from n = 317 ROIs, aligned to mice’s first lick during the sound (black) or answer (magenta) period, denoted by the gray dotted line. **D)** Summary PSTHs for Ca^2+^ events in n = 317 ROIs, aligned to the first lick (gray dotted line) during the sound or answer periods (black and magenta, respectively). Shading in C and D is ± SEM.

We summarized these results by comparing latency histograms of dendritic spike times aligned to the first lick during the sound and answer period (33 ms bin-widths; Figure 9D). A two-way, repeated measures ANOVA revealed a main effect of lick alignment mode (e.g., aligned to first lick in sound or answer; F(1,1316) = 192.7, p<0.0001), latency bin (F(36,11376) = 10.34, p<0.0001), and a latency bin x alignment mode interaction (F(36,11376) = 11.52, p<0.0001). Altogether, these data show that licking during sound presentation is not correlated with dendritic electrogenesis, thereby providing a mechanistic explanation for how increases in dendritic event rates are limited to the answer period in our task.

### Sound presentation modulates dendritic spike amplitude

The peak amplitude of dendritic spikes in our datasets varied over ∼10 fold (Figure 2). Thus, dendritic spikes are unlikely to be all-or-none events, but rather could control axo-somatic burst firing in a graded manner. Does dendritic spike amplitude vary systematically during specific behaviorally relevant trial epochs? We tested this idea by comparing dendritic spike amplitudes during the sound or answer period with those occurring randomly during a 2-3 s baseline period prior to sound onset of each trial. Sound excited dendrites showed a ∼40 fold increase in dendritic spike probability and a 22% increase in dendritic spike amplitude relative to the baseline period (Figure 10A-C. Median peak amplitude in baseline and sound: 2.8 vs 3.34 ΛF/F, p = 0.008, signed-rank test). By contrast, sound inhibited dendrites showed a profound sound-evoked reduction in dendritic spike probability, and the remaining dendritic spikes occurring during sound presentation were of significantly smaller amplitude than those in the trial baseline (Figure 10D-F. Median peak amplitude in baseline and sound: 3.9 vs. 3.3 ΛF/F, p<0.0001, signed-rank test). Interestingly, neither early nor late answer excited dendrites showed a significant change in dendritic spike amplitude during the answer period, despite the profound increase in event probability relative to baseline (Figure 10G-L). Altogether these results indicate that although dendritic spike probability increases during the sound and answer period, the amplitude of dendritic spikes is selectively and bi-directionally modulated during sound presentation.

**Figure 10:**
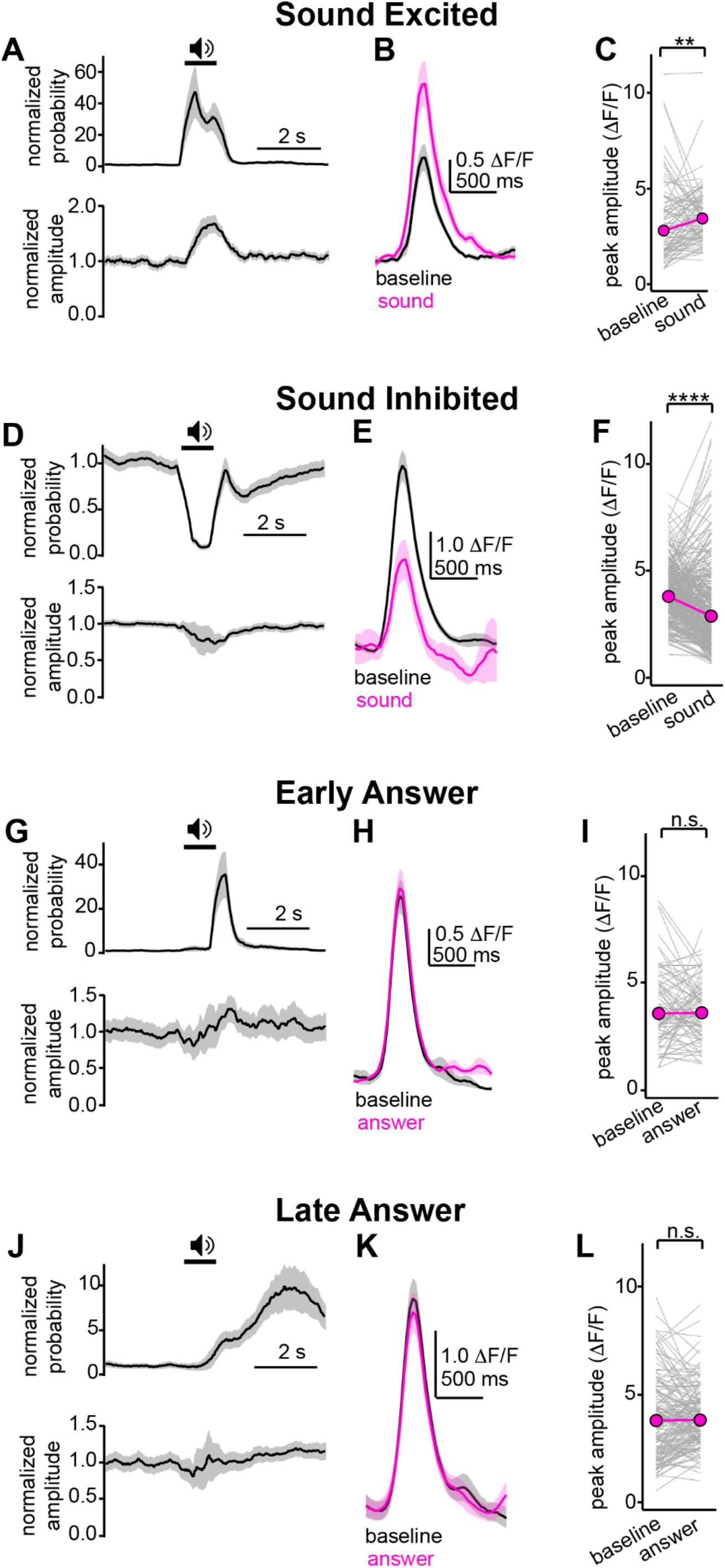
Dendritic spike magnitudes are modulated by sound, but not answer period. **A)** Normalized event probability and peak amplitude (upper and lower panels, respectively) plotted across time for all sound excited dendrites. **B)** Example averaged events in a single sound excited dendrite in baseline (black) and sound (magenta) periods. Of note is the larger event amplitude during sound presentation. In tandem with data from panel A, these results indicate that sound presentation significantly enhances dendritic spike amplitudes in sound excited neurons. **C)** Summary of Ca^2+^ event peak amplitudes in baseline and during sound presentation for n = 91 sound excited dendrites. Gray lines are individual dendrites, magenta is median. **D-F)** Same as A-C, but for sound inhibited dendrites. Of note, sound presentation drastically reduces dendritic spike probability, and the few events remaining during sound presentation have significantly reduced amplitudes. **G-L)** Same as A-C; D-F, but for early (G-I) and late (J-L) answer modulated dendrites. Although event rates increase during the answer period in these dendrites, peak amplitudes of individual Ca^2+^ events are not significantly different between the baseline and answer periods (early answer: 3.59 vs 3.62 Λ1F/F, p = 0.44. Late answer: 3.79 vs 3.81, p=0.28 for baseline and answer period, signed-rank test). ** = p < 0.01, **** = p<0.0001.

### Silencing answer period activity impairs discriminative learning

Corticofugal neurons are important for many learned and innate behaviors (Bajo et al., 2010; Znamenskiy and Zador, 2013; Liu et al., 2016; Ruediger and Scanziani, 2020; Tang and Higley, 2020), and dendritic spikes could be potential effectors of synaptic plasticity to support these processes (Ranganathan et al., 2018). We find that a major fraction of dendritic spikes occur in the answer period *after* the discriminative sound has ended (Figures 3-6), and that this activity is observable even in untrained and non-task engaged mice (Figure 8). Does this answer period activity contribute to the learning-related functions of corticofugal neurons? If true, selective silencing of auditory corticofugal neurons *after* sound presentation should suffice to impair learning.

To this end, we bilaterally expressed a soma-targeted version of the inhibitory opsin GtACR1 in auditory corticofugal neurons by injecting a retrograde-cre recombinase virus in the IC of GtACR1 fl/fl mice (Figure 11A; see also Li et al., 2019). We first confirmed that GtACR1 silenced auditory corticofugal activity using patch-clamp recordings in acute auditory cortex slices from injected GtACR1 fl/fl mice. Activating GtACR1 via 625 nm light hyperpolarized auditory corticofugal neurons and potently reduced neuronal excitability, such that >3 mW peak power largely abolished spiking even to unphysiologically strong current steps (Figure 9B,C). Indeed, LED power even as low as 0.5 mW sufficed to significantly dampen firing rate-intensity curves in corticofugal neurons (Figure 11D, n = 9 cells from 2 mice. Main effects of current step F(5,40)=89.62, p<0.0001 + LED activation F(1,8)=71.63, p<0.0001, and current step x LED activation interaction F(5,28)=6.494, p=0.0004, mixed-effects model), and we never observed rebound spiking due to shifts in Cl-reversal potential under our conditions. These data show that GtACR1 activation strongly dampens corticofugal neuron excitability.

**Figure 11:**
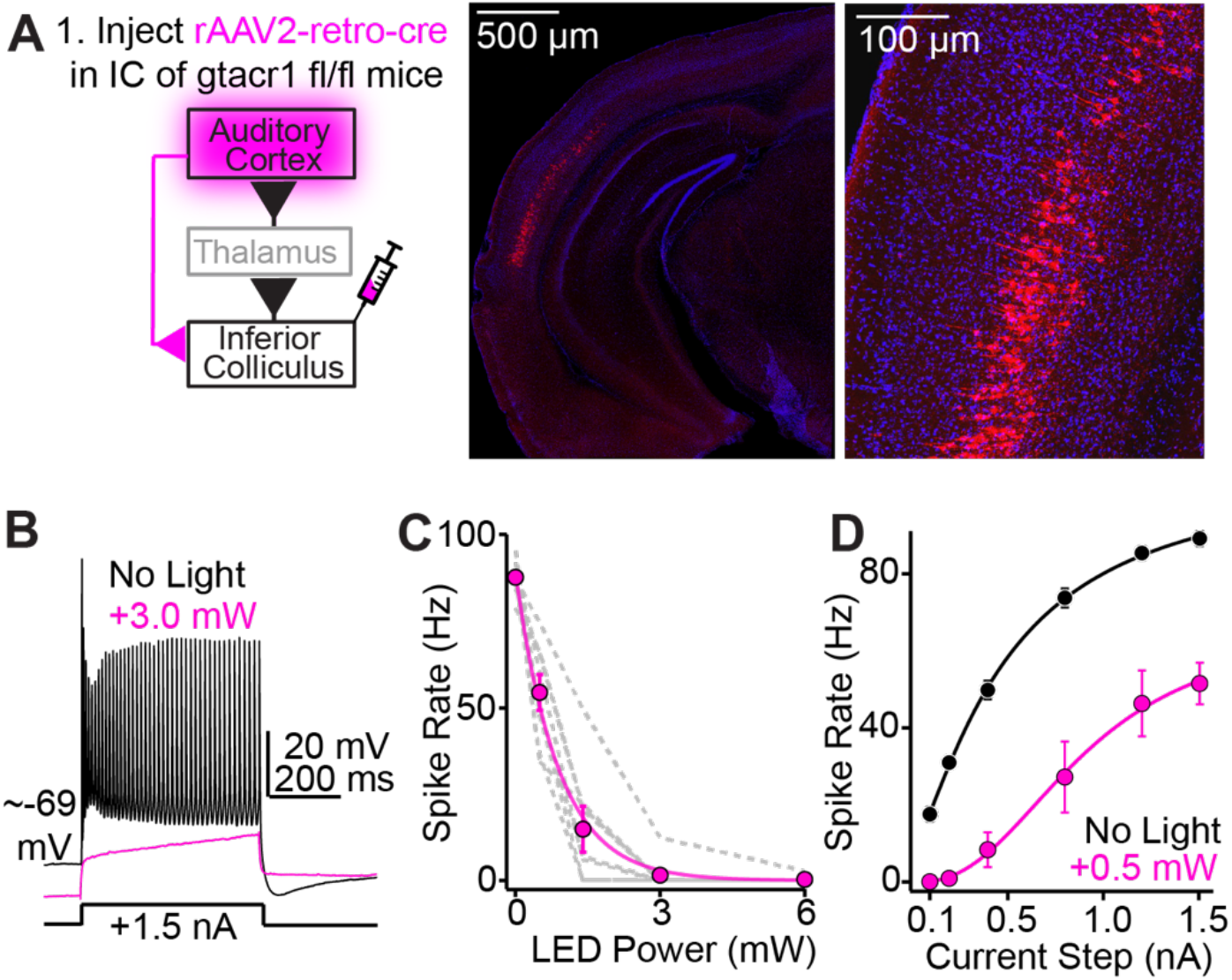
GtACR1 activation silences auditory corticofugal neurons. **A)** Cartoon of experiment. Retrograde-cre is injected in the IC of GtACR1 fl/fl mice. Whole-cell patch clamp recordings are obtained ∼2 weeks later from GtACR1 expressing corticofugal layer 5 neurons in acute auditory cortex brain slices. GtACR1 is activated via a 625 nm coupled optic fiber positioned above the slice. Right: Micrographs of GtACR1 expression in layer 5 corticofugal neurons of auditory cortex. The FusionRed tag, and thus GtACR1 construct, is soma-targeted in these mice (Li et al., 2019). **B)** Example recording showing spike responses to 1.5 nA current steps in absence or presence of GtACR1 activation (3 mW; black and magenta traces, respectively). **C)** Summary showing the effect of increasing light levels on spike rates during 1.5 nA current steps (n=8 cells from N = 2 mice). Of note is the near complete absence of spiking with >3 mW light. Gray lines are individual neurons, magenta points are mean ± SEM. Magenta line is mono-exponential fit to mean data. **D)** Summary data of firing rate intensity curves in absence or presence of GtACR1 activation (n = 9 cells from N = 2 mice).

We next tested if silencing corticofugal neurons in the answer period impacts mice’s learning of the GO/NO-GO sAM task by comparing the learning rates of GtACR1 fl/fl mice injected with a retrograde-cre virus or control saline buffer (Figure 12A). Mice were trained using a standardized protocol, in which auditory corticofugal neurons were silenced bilaterally via light delivered to the auditory cortex *only* during the answer period on each trial (Figure 12B).

**Figure 12:**
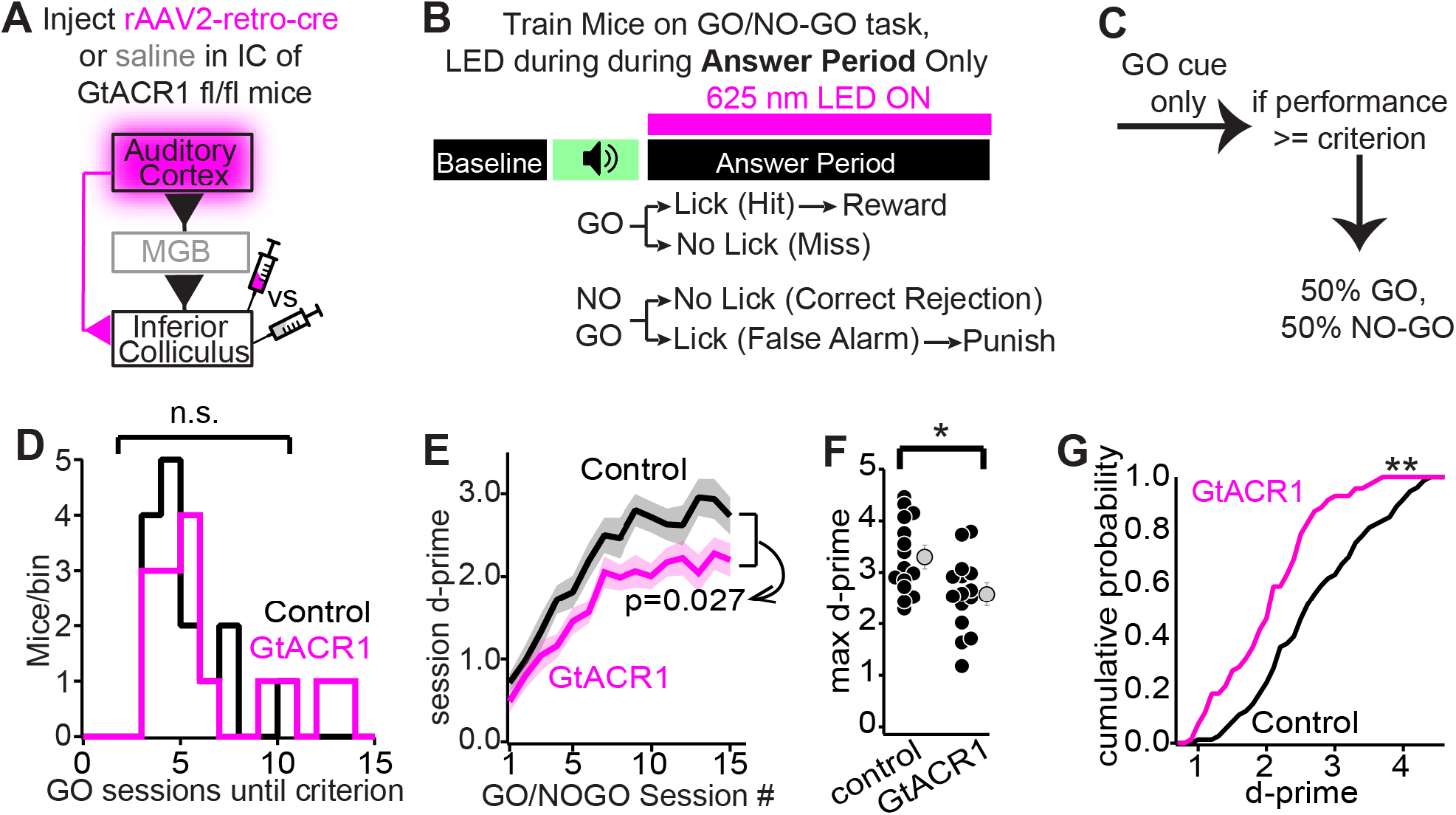
Silencing auditory corticofugal neurons in the answer period impairs discriminative learning. **A)** A retrograde-cre virus or control buffer is injected bilaterally in the IC of GtACR1 fl/fl mice. **B)** Mice are trained on the GO/NO-GO discrimination task. Bilateral LEDs over auditory cortices are activated on all trials *only* during the answer period. **C)** Mice are initially trained to associate the GO sound with rewards, and upon reaching criterion, are trained on the full GO/NO-GO task. **D)** Histograms of the number of GO-only sessions until criterion for control (black) and GtACR1 mice (magenta) **E)** Mice’s d-prime across GO/NO-GO training sessions. Lines + shading are mean ± SEM of each group **F)** Maximum d-prime across sessions 11-15 for both groups of mice. Black dots are individual mice, grey is mean ± SEM. **G)** Cumulative distribution of d-prime values in sessions 11-15 for control and GtACR1 expressing mice. * = p < 0.05, ** = p < 0.01.

Because GtACR1 is not activated during the sound, any effect should be independent of a corticofugal contribution to perception or discrimination. Mice were first trained to associate a GO sound with reward availability, and upon reaching criterion (>75% hits on two consecutive sessions), trained to discriminate GO and NO-GO sounds presented with 50% probability (Figure 12C). Both mouse groups showed similar increases in hit rates when exposed only to GO cues (Figure 12D. Median number of sessions to criterion: 4 and 5 for control and GtACR1 mice, respectively. p=0.89, Kolmogorov-Smirnov test). Thus, silencing corticofugal neurons in the answer period does not appear to impair mice’s motivation or ability to learn simple, operant associations. By contrast, GtACR1-expressing mice showed a significant reduction in d-prime across GO/NO-GO training sessions compared to control mice (Figure 12E; mixed effects model. Main effects of session and treatment group: F(6.44,178.1)=63.42, p<0.0001 and F(1,28)=5.479, p=0.027, respectively), and the maximum d-prime reached across all sessions was significantly lower for GtACR1 compared to control mice (Figure 12F; 3.3 ± 0.2 vs 2.6 ± 0.2 for n = 15 control and n = 15 GtACR1 expressing mice, t(28)=2.74, p=0.010, unpaired t-test). Consequently, the cumulative distribution of d-prime values in sessions 11-15, during which mice’s discrimination has typically reached a plateau (Figure 12E; see also Figure 1E), significantly differed between control and GtACR1 mice (Figure 12G. K-S statistic = 0.299, p=0.002, Kolmogorov-Smirnov test). These group-level differences do not reflect impaired learning from cre recombinase or GtACR1 expression in corticofugal neurons, as a separate cohort of n = 11 control and n = 12 GtACR1-expressing mice trained in absence of LED activation showed similar sensitivity indices across sessions (Mixed-effects model. Main effect of Session #: F(4.553, 95.61)=22.46, p<0.001; No effect of treatment group: F(1,21)=0.8053, p=0.38).

### Silencing corticofugal neurons does not impair mice’s licking

The results of experiments in Figure 12 could reflect learning-related deficits. Alternatively these results could arise if corticofugal silencing impacts mice’s licking patterns. To disambiguate these two scenarios, we analyzed control and virus expressing mice’s licking patterns across learning. Lick histograms averaged across all sessions were similar in the two mouse groups, and showed no appreciable change upon light presentation on either hit or false alarm trials (Figure 13A). We also compared control and virus mice’s absolute number of licks (Figure 13B) and first lick latency (Figures 13C,D) during the answer period of hit and false alarm trials in each GO and NO-GO session. Mixed-effects models revealed a main effect of session number on absolute lick count for hit (F(5.21,144)=2.29, p=0.046) and false alarm trials (F(6.58,182)=6.29, p<0.0001), indicating a modest, learning-dependent decrease in mice’s licks in the answer period. However, there was no main effect of treatment group (F(1,28)=0.0003, p=0.98 and F(1,28)=0.20, p=0.66 for hit and false alarm trials, respectively), indicating that this decrease was similar in control and virus mice (Figure 13B). Additionally, there was no main effect of session number or treatment group on mice’s first lick latency in the answer period for hit (Figure 7C. F(3.85,106.60)=1.032, p=0.39; F(1,28)=0.37, p=0.54 for session number and treatment group, respectively) and false alarm trials (Figure 7D. F(4.97,137.6)=0.6161, p=0.69; F(1,28)=0.58, p=0.45 for session number and treatment group, respectively). We conclude that optogenetic silencing during the answer period impacts discriminative learning rather than the initiation or execution of licking, thereby intimating an unexpected non-sensory component to corticofugal function.

**Figure 13:**
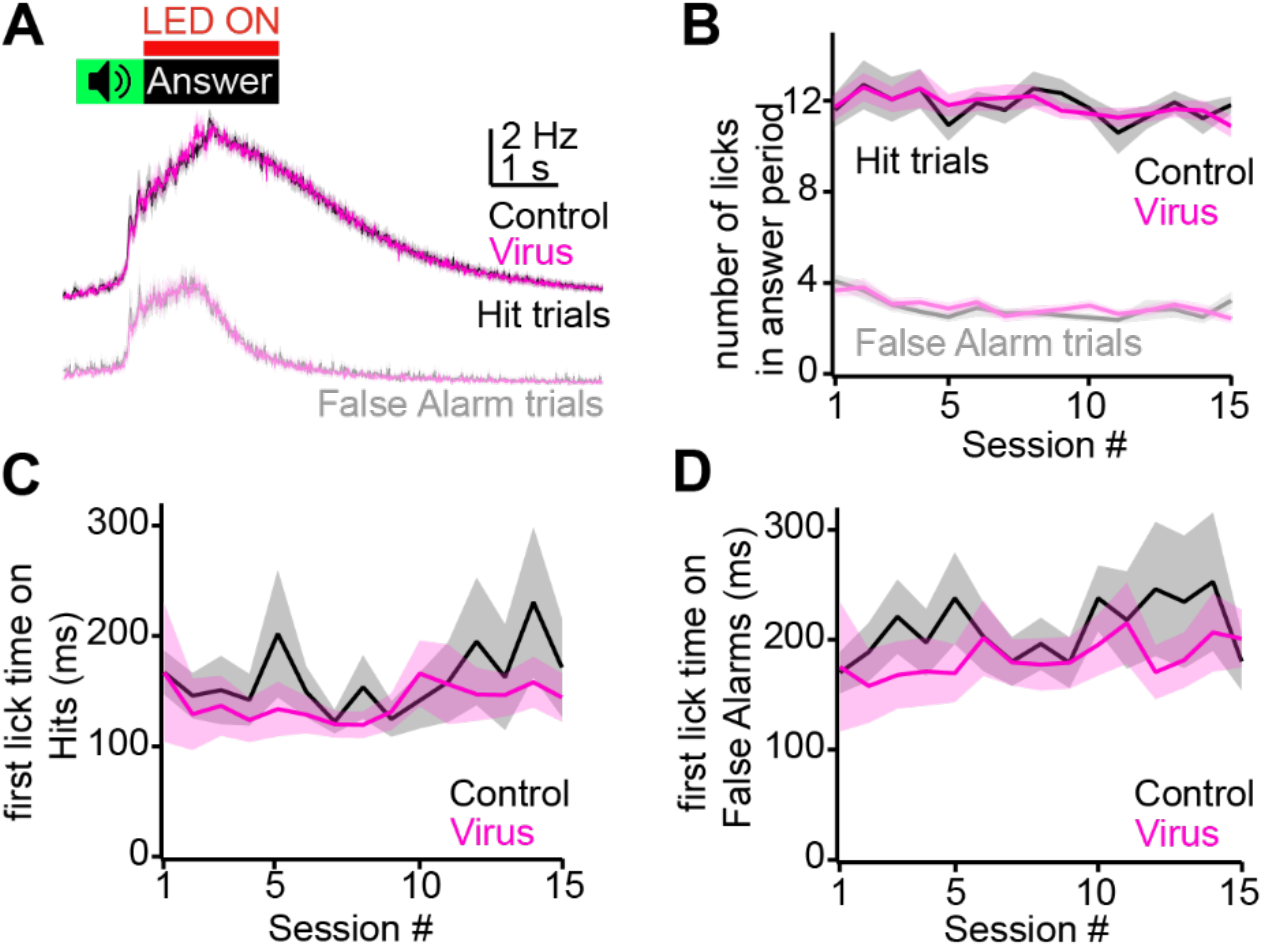
Optogenetic silencing in the answer period does not impair mice’s licking behavior or reaction times. **A)** Lick histograms across averaged all behavioral sessions for n = 15 control and n = 15 virus mice (black and magenta, respectively), on hit and false alarm trials (upper and lower panels, respectively). Green denotes sound presentation, red line denotes optogenetic LED activation in the answer period. **B)** Summary data showing the number of licks in the answer period of hit and false alarm trials for each session, in control and virus mice. **C,D)** Latency of first lick in the answer period following sound offset is plotted for control and virus mice on hit (C) and false alarm (D) trials. Color scheme in B-D is same as in A. Shading for all graphs is ± SEM.

## Discussion

By switching neuronal spiking from low-frequency trains to high-frequency bursts, dendritic Ca^2+^ spikes trigger a uniquely strong output signal (Xu et al., 2012a; Takahashi et al., 2016, 2020; Ranganathan et al., 2018; Beaulieu-Laroche et al., 2019; Francioni et al., 2019; Trautmann et al., 2021) reflecting correlated activity in basal and apical dendrites (Larkum et al., 1999, 2022; Takahashi and Magee, 2009; Bittner et al., 2015). In auditory corticofugal neurons of behaving mice, the majority of dendritic spikes occurred *after* the discriminative sound stimulus and correlated with mice’s purposeful actions during the trial answer period. Silencing corticofugal neurons during the answer period impaired discriminative learning, highlighting a novel function for the abundance of non-sensory signals observed throughout sensory cortices (Musall et al., 2019; Stringer et al., 2019). An important consideration is that this manipulation slowed, but did not abolish learning; answer period activity may be one of many mechanisms that contribute to learning under our conditions. We also may not have silenced all auditory corticofugal neurons, since our retrograde virus does not transduce layer 6 neurons (Tervo et al., 2016) which contribute 10-20% of the auditory cortico-collicular pathway (Yudnitsev et al., 2022). Rather, our results place a lower bound on auditory corticofugal neurons’ contribution to learning.

Answer period activity co-varied with mice’s licking and may reflect motor-related inputs from secondary motor cortex, basal forebrain, or basal ganglia synapsing onto apical tuft dendrites (Schneider et al., 2014; Nelson and Mooney, 2016; Clayton et al., 2021). These projections could transmit motor efference copies or potentially, a learned representation of rewarded actions (Peters et al., 2017; Park et al., 2022). Answer period activity could also reflect force exertion, as recently suggested for midbrain dopamine neurons (Bakhurin et al., 2023).

Although a motor-related origin is perhaps the simplest interpretation, we cannot rule out that answer period activity reflects an arousal signal that correlates with, but is not directly driven by mice’s action initiation. Indeed, dendritic spikes arise via R-type Ca^2+^ channels (Takahashi and Magee, 2009; Williams and Fletcher, 2019), with potential contributions from NMDA receptors (Larkum et al., 2009). Both NMDA receptors and R-Type channels are potently regulated by neuromodulators such as noradrenaline and acetylcholine (Raman et al., 1996; Williams and Fletcher, 2019) that impact dendritic electrogenesis (Labarrera et al., 2018; Williams and Fletcher, 2019). However, dendritic activity is yoked to the answer period rather than anticipatory licks during sound presentation, which should mark the onset of any arousal increase (Figure 9). Thus, a neuromodulatory origin may be less likely given these temporal dynamics. Future studies are necessary to dissect disentangle the mechanistic bases of this non-auditory activity.

In somatosensory cortex, dendritic spikes are triggered by reward consumption, with the reliability of this reward-evoked activity increasing across learning (Lacefield et al., 2019). Our results contrast somewhat with Lacefield et al, as dendritic activity occurred regardless of reward consumption. However, this distinction may reflect different task designs: The rewarded action in the Lacefield study was manual lever release, whereas licking during the answer period was operantly reinforced in our task. Alternatively, these differences may reflect unique properties of auditory and somatosensory cortices. We also cannot rule out any learning-dependent changes in answer period activity (Takamiya et al., 2022); future studies will be required to address this knowledge gap. Nevertheless, dendritic spikes can occur in absence of our task engagement, suggesting that these signals are present across the learning process.

### Corticofugal dendritic activity is often inhibited by sounds

Spontaneous dendritic activity was often inhibited by sound presentation. Dendritic spike amplitudes were also reduced, suggesting a role for postsynaptic inhibition (Pérez-Garci et al., 2006) rather than a simple reduction in the rate of presynaptic excitation. Although this “inhibition” may reflect an increased excitation that depolarizes the distal dendrites and inactivates Ca^2+^ channels as in thalamic relay neurons (Zhan et al., 2000), channel inactivation cannot explain our results: Dendritic spikes in pyramidal neurons are mediated by R-type Ca^2+^ channels rather than the low-voltage inactivated T-type channels of thalamic relay neurons (Takahashi and Magee, 2009; Williams and Fletcher, 2019); the amplitude and duration of dendritic spikes thus increases, rather than decreases, with prior depolarizations (Xu et al., 2012a; Harnett et al., 2013). Our results are instead more consistent with sound presentation recruiting dendrite targeting inhibitory neurons such as Martinotti cells (Hilscher et al., 2017) or layer 1 interneurons (Takesian et al., 2018).

### Dendritic Ca^2+^ spikes as effectors of supervised plasticity

Our data further dovetail with studies intimating a learning-related role for layer 5 neurons of sensory cortex (Bajo et al., 2010; Ruediger and Scanziani, 2020). A tacit assumption is that learning impairments stemming from layer 5 pyramidal neuron silencing reflect a loss of sensory-evoked corticofugal signals, which could support learning by increasing sensory discrimination (Znamenskiy and Zador, 2013). Our data now indicate a contribution of non-auditory signals transmitted several seconds after a discriminative stimulus has ended, in agreement with data showing that “trial outcome” or “action timing” activity contributes to sensorimotor learning (Namboodiri et al., 2015; Levy et al., 2020; Schoenfeld et al., 2021).

How might non-auditory activity support learning? One hypothesis is that dendritic spikes in pyramidal neurons may act be cellular correlates of model-free “supervised” learning algorithms (Sutton and Barto, 2018), similar to the hypothesized function of dendritic Ca^2+^ spikes in the Purkinje and Purkinje-like cells of mammalian cerebellum and mormyrid fish’s electrosensory lobe (Bell et al., 2008; Raymond and Medina, 2018). In hippocampus CA1 neurons, dendritic Ca^2+^ spikes potentiate excitatory synapses active even >1 s prior in time (Bittner et al., 2017), thereby instructing plasticity at synapses whose activity precedes, but does not directly initiate dendritic electrogenesis. If similar mechanisms exist in layer 5 neurons, answer period activity could promote heterosynaptic potentiation of “bottom-up” synapses transmitting information regarding the discriminative sound stimuli. Such a plasticity rule could account for the biased selectivity of sound-excited dendrites towards GO cues (Figure 3), and might also explain classic findings of sound feature over-representation during development (Zhang et al., 2001), Pavlovian conditioning (Edeline et al., 1993), and instrumental learning (Polley et al., 2006; Bieszczad and Weinberger, 2010). A second, mutually compatible hypothesis is that the action potential bursts triggered by dendritic spikes instruct plasticity in target brain regions. Auditory corticofugal neurons are famously required for re-learning of a sound localization task upon hearing loss (Bajo et al., 2010), but apparently dispensable for task performance after re-learning (Bajo et al., 2019). Thus learning-related plasticity could occur in postsynaptic targets of corticofugal neurons. In agreement, auditory cortex orchestrates receptive field plasticity of mammalian IC neurons (Yan and Suga, 1998; Suga et al., 2002; Ji et al., 2005), whereas avian IC neurons exhibit learning- and NMDA receptor-dependent plasticity (Brainard and Knudsen, 1993; Feldman et al., 1996; Penzo and Peña, 2009). Interestingly, auditory corticofugal activity promotes cooperative NMDA receptor activation at ascending synapses in IC neurons (Oberle et al., 2022). If IC neurons are capable of long timescale learning rules, corticofugal neuron burst firing could promote plasticity at “bottom-up” synapses in early brain regions. Alternatively, learning-related plasticity could be independent of midbrain circuits, given the brain-wide collateralization of corticofugal neurons (Chen et al., 2019; Williamson and Polley, 2019); target-specific silencing of axonal collaterals could shed light on this knowledge gap (Copits et al., 2021; Mahn et al., 2021).

Three caveats of our optogenetic approach must be highlighted. First, this manipulation silences both action potential output and dendritic Ca^2+^ spikes. Since backpropagating action potentials are necessary for dendritic electrogenesis (Larkum et al., 1999; Takahashi and Magee, 2009), we cannot disambiguate if silencing corticofugal neurons impairs learning by removing a cell-intrinsic or efferent signal. Second, we cannot determine if burst firing, rather than an equivalent number of low frequency simple spikes, is explicitly necessary for learning-related functions of corticofugal neurons. Third, our approach not only silences answer period activity, but potentially also the spontaneous spiking of opsin expressing, but non-task responsive corticofugal neurons. Removal of this background activity could in principle contribute to our observed learning deficits, although we think this possibility unlikely: Inhibiting corticofugal neurons under acoustically silent conditions does not impact spontaneous firing rates of IC neurons (Blackwell et al., 2020), suggesting that spontaneous spikes contribute minimally to downstream network function.

Non-auditory activity is also abundant in the IC of awake animals (Metzger et al., 2006; Yang et al., 2020; Shaheen et al., 2021; Quass et al., 2023), and seemingly persists in absence of corticofugal feedback (Lee et al., 2023). Thus it is currently unclear if non-auditory activity in corticofugal neurons is the cause, or rather the consequence of similar activity patterns in IC neurons. Alternatively, non-auditory activity could reflect reverberatory interactions between brain regions, akin to the ”ramping” signals of thalamo-cortico-thalamic motor pathways (Guo et al., 2017). Indeed, auditory cortex neurons persistently fire during delay periods of working memory tasks (Gottlieb et al., 1989; Huang et al., 2016; but see Yu et al., 2021), with similar activity observable in IC (Metzger et al., 2006). Thus, the auditory system could maintain non-auditory activity via dynamic interactions between evolutionarily disparate circuits.

## Acknowledgements

We thank Drs. Kevin Bender, Travis Moschak, and Gideon Rothschild for thoughtful comments on the manuscript, and Mr. Deepak Dileepkumar for constructing the sound-attenuating chambers + optogenetic LEDs. Funding was generously provided by the Hearing Health Foundation, the Whitehall Foundation, and NIH grant R01DC019090 (PFA).

## Notes

### Competing Interest Statement

The authors have declared no competing interest.

### Summary of Updates

Major revisions

## References

Apostolides PF, Milstein AD, Grienberger C, Bittner KC, Magee JC (2016) Axonal Filtering Allows Reliable Output during Dendritic Plateau-Driven Complex Spiking in CA1 Neurons. Neuron 89:770–783.

Bajo VM, Nodal FR, Korn C, Constantinescu AO, Mann EO, Boyden ES, King AJ (2019) Silencing cortical activity during sound-localization training impairs auditory perceptual learning. Nat Commun 10:3075.

Bajo VM, Nodal FR, Moore DR, King AJ (2010) The descending corticocollicular pathway mediates learning-induced auditory plasticity. Nat Neurosci 13:253–260.

Bakhurin K, Hughes RN, Jiang Q, Hossain M, Gutkin B, Fallon IP, Yin HH (2023) Force tuning explains changes in phasic dopamine signaling during stimulus-reward learning. Neuroscience. Available at: http://biorxiv.org/lookup/doi/10.1101/2023.04.23.537994 [Accessed November 1, 2023].

Bandyopadhyay S, Shamma SA, Kanold PO (2010) Dichotomy of functional organization in the mouse auditory cortex. Nat Neurosci 13:361–368.

Banerjee A, Seriès P, Pouget A (2008) Dynamical constraints on using precise spike timing to compute in recurrent cortical networks. Neural Comput 20:974–993.

Beaulieu-Laroche L, Toloza EHS, Brown NJ, Harnett MT (2019) Widespread and Highly Correlated Somato-dendritic Activity in Cortical Layer 5 Neurons. Neuron 103:235–241.e4.

Bell CC, Han V, Sawtell NB (2008) Cerebellum-like structures and their implications for cerebellar function. Annu Rev Neurosci 31:1–24.

Bieszczad KM, Weinberger NM (2010) Representational gain in cortical area underlies increase of memory strength. Proc Natl Acad Sci U S A 107:3793–3798.

Bittner KC, Grienberger C, Vaidya SP, Milstein AD, Macklin JJ, Suh J, Tonegawa S, Magee JC (2015) Conjunctive input processing drives feature selectivity in hippocampal CA1 neurons. Nat Neurosci 18:1133–1142.

Bittner KC, Milstein AD, Grienberger C, Romani S, Magee JC (2017) Behavioral time scale synaptic plasticity underlies CA1 place fields. Science 357:1033–1036.

Brainard M, Knudsen E (1993) Experience-dependent plasticity in the inferior colliculus: a site for visual calibration of the neural representation of auditory space in the barn owl. J Neurosci 13:4589– 4608.

Chen X, Sun Y-C, Zhan H, Kebschull JM, Fischer S, Matho K, Huang ZJ, Gillis J, Zador AM (2019) High-Throughput Mapping of Long-Range Neuronal Projection Using In Situ Sequencing. Cell 179:772–786.e19.

Clark BA, Cull-Candy SG (2002) Activity-Dependent Recruitment of Extrasynaptic NMDA Receptor Activation at an AMPA Receptor-Only Synapse. J Neurosci 22:4428–4436.

Clayton KK, Williamson RS, Hancock KE, Tasaka G-I, Mizrahi A, Hackett TA, Polley DB (2021) Auditory Corticothalamic Neurons Are Recruited by Motor Preparatory Inputs. Curr Biol CB 31:310–321.e5.

Copits BA et al. (2021) A photoswitchable GPCR-based opsin for presynaptic inhibition. Neuron 109:1791–1809.e11.

Doron G, Shin JN, Takahashi N, Drüke M, Bocklisch C, Skenderi S, de Mont L, Toumazou M, Ledderose J, Brecht M, Naud R, Larkum ME (2020) Perirhinal input to neocortical layer 1 controls learning. Science 370:eaaz3136.

Edeline JM, Pham P, Weinberger NM (1993) Rapid development of learning-induced receptive field plasticity in the auditory cortex. Behav Neurosci 107:539–551.

Feldman DE, Brainard MS, Knudsen EI (1996) Newly learned auditory responses mediated by NMDA receptors in the owl inferior colliculus. Science 271:525–528.

Fletcher LN, Williams SR (2019) Neocortical Topology Governs the Dendritic Integrative Capacity of Layer 5 Pyramidal Neurons. Neuron 101:76–90.e4.

Francioni V, Padamsey Z, Rochefort NL (2019) High and asymmetric somato-dendritic coupling of V1 layer 5 neurons independent of visual stimulation and locomotion. eLife 8:e49145.

Galloni AR, Laffere A, Rancz E (2020) Apical length governs computational diversity of layer 5 pyramidal neurons. eLife 9.

Gottlieb Y, Vaadia E, Abeles M (1989) Single unit activity in the auditory cortex of a monkey performing a short term memory task. Exp Brain Res 74:139–148.

Guo ZV, Inagaki HK, Daie K, Druckmann S, Gerfen CR, Svoboda K (2017) Maintenance of persistent activity in a frontal thalamocortical loop. Nature 545:181–186.

Harnett MT, Xu N-L, Magee JC, Williams SR (2013) Potassium Channels Control the Interaction between Active Dendritic Integration Compartments in Layer 5 Cortical Pyramidal Neurons. Neuron 79:516–529.

Hilscher MM, Leão RN, Edwards SJ, Leão KE, Kullander K (2017) Chrna2-Martinotti Cells Synchronize Layer 5 Type A Pyramidal Cells via Rebound Excitation. PLoS Biol 15:e2001392.

Hires SA, Zhu Y, Tsien RY (2008) Optical measurement of synaptic glutamate spillover and reuptake by linker optimized glutamate-sensitive fluorescent reporters. Proc Natl Acad Sci 105:4411–4416.

Huang Y, Heil P, Brosch M (2019) Associations between sounds and actions in early auditory cortex of nonhuman primates. eLife 8:e43281.

Huang Y, Matysiak A, Heil P, König R, Brosch M (2016) Persistent neural activity in auditory cortex is related to auditory working memory in humans and nonhuman primates. eLife 5:e15441.

Hubel DH, Wiesel TN (1963) Shape and arrangement of columns in cat’s striate cortex. J Physiol 165:559–568.

Issa JB, Haeffele BD, Agarwal A, Bergles DE, Young ED, Yue DT (2014) Multiscale optical Ca2+ imaging of tonal organization in mouse auditory cortex. Neuron 83:944–959.

Ji W, Suga N, Gao E (2005) Effects of agonists and antagonists of NMDA and ACh receptors on plasticity of bat auditory system elicited by fear conditioning. J Neurophysiol 94:1199–1211.

Joshi A, Middleton JW, Anderson CT, Borges K, Suter BA, Shepherd GMG, Tzounopoulos T (2015) Cell-specific activity-dependent fractionation of layer 2/3→5B excitatory signaling in mouse auditory cortex. J Neurosci Off J Soc Neurosci 35:3112–3123.

Kerlin A, Mohar B, Flickinger D, MacLennan BJ, Dean MB, Davis C, Spruston N, Svoboda K (2019) Functional clustering of dendritic activity during decision-making. eLife 8.

Kim U, McCormick DA (1998) The Functional Influence of Burst and Tonic Firing Mode on Synaptic Interactions in the Thalamus. J Neurosci 18:9500–9516.

Kreitzer AC, Regehr WG (2000) Modulation of Transmission during Trains at a Cerebellar Synapse. J Neurosci 20:1348–1357.

Labarrera C, Deitcher Y, Dudai A, Weiner B, Kaduri Amichai A, Zylbermann N, London M (2018) Adrenergic Modulation Regulates the Dendritic Excitability of Layer 5 Pyramidal Neurons In Vivo. Cell Rep 23:1034–1044.

Lacefield CO, Pnevmatikakis EA, Paninski L, Bruno RM (2019) Reinforcement Learning Recruits Somata and Apical Dendrites across Layers of Primary Sensory Cortex. Cell Rep 26:2000–2008.e2.

Larkum ME, Nevian T, Sandler M, Polsky A, Schiller J (2009) Synaptic integration in tuft dendrites of layer 5 pyramidal neurons: a new unifying principle. Science 325:756–760.

Larkum ME, Wu J, Duverdin SA, Gidon A (2022) The Guide to Dendritic Spikes of the Mammalian Cortex In Vitro and In Vivo. Neuroscience 489:15–33.

Larkum ME, Zhu JJ, Sakmann B (1999) A new cellular mechanism for coupling inputs arriving at different cortical layers. Nature 398:338–341.

Lee J, Rothschild G (2021) Encoding of acquired sound-sequence salience by auditory cortical offset responses. Cell Rep 37:109927.

Lee T-Y, Weissenberger Y, King AJ, Dahmen JC (2023) Midbrain encodes sound detection behavior without auditory cortex. Neuroscience. Available at: http://biorxiv.org/lookup/doi/10.1101/2023.06.07.544013 [Accessed June 27, 2023].

Lesicko AMH, Angeloni CF, Blackwell JM, De Biasi M, Geffen MN (2022) Corticofugal regulation of predictive coding. eLife 11:e73289.

Levy S, Lavzin M, Benisty H, Ghanayim A, Dubin U, Achvat S, Brosh Z, Aeed F, Mensh BD, Schiller Y, Meir R, Barak O, Talmon R, Hantman AW, Schiller J (2020) Cell-Type-Specific Outcome Representation in the Primary Motor Cortex. Neuron 107:954–971.e9.

Li N, Chen S, Guo ZV, Chen H, Huo Y, Inagaki HK, Chen G, Davis C, Hansel D, Guo C, Svoboda K (2019) Spatiotemporal constraints on optogenetic inactivation in cortical circuits. eLife 8:e48622.

Lisman J (1997) Bursts as a unit of neural information: making unreliable synapses reliable. Trends Neurosci 20:38–43.

Liu B-H, Huberman AD, Scanziani M (2016) Cortico-fugal output from visual cortex promotes plasticity of innate motor behaviour. Nature 538:383–387.

Liu J, Whiteway MR, Sheikhattar A, Butts DA, Babadi B, Kanold PO (2019) Parallel Processing of Sound Dynamics across Mouse Auditory Cortex via Spatially Patterned Thalamic Inputs and Distinct Areal Intracortical Circuits. Cell Rep 27:872–885.e7.

London M, Roth A, Beeren L, Häusser M, Latham PE (2010) Sensitivity to perturbations in vivo implies high noise and suggests rate coding in cortex. Nature 466:123–127.

Mahn M et al. (2021) Efficient optogenetic silencing of neurotransmitter release with a mosquito rhodopsin. Neuron 109:1621–1635.e8.

Merzenich MM, Knight PL, Roth GL (1975) Representation of cochlea within primary auditory cortex in the cat. J Neurophysiol 38:231–249.

Metzger RR, Greene NT, Porter KK, Groh JM (2006) Effects of reward and behavioral context on neural activity in the primate inferior colliculus. J Neurosci Off J Soc Neurosci 26:7468–7476.

Morest DK, Oliver DL (1984) The neuronal architecture of the inferior colliculus in the cat: defining the functional anatomy of the auditory midbrain. J Comp Neurol 222:209–236.

Musall S, Kaufman MT, Juavinett AL, Gluf S, Churchland AK (2019) Single-trial neural dynamics are dominated by richly varied movements. Nat Neurosci 22:1677–1686.

Nahir B, Jahr CE (2013) Activation of Extrasynaptic NMDARs at Individual Parallel Fiber–Molecular Layer Interneuron Synapses in Cerebellum. J Neurosci 33:16323–16333.

Namboodiri VMK, Huertas MA, Monk KJ, Shouval HZ, Hussain Shuler MG (2015) Visually cued action timing in the primary visual cortex. Neuron 86:319–330.

Naud R, Sprekeler H (2018) Sparse bursts optimize information transmission in a multiplexed neural code. Proc Natl Acad Sci 115:E6329–E6338.

Nelson A, Mooney R (2016) The Basal Forebrain and Motor Cortex Provide Convergent yet Distinct Movement-Related Inputs to the Auditory Cortex. Neuron 90:635–648.

Neubrandt M, Oláh VJ, Brunner J, Marosi EL, Soltesz I, Szabadics J (2018) Single Bursts of Individual Granule Cells Functionally Rearrange Feedforward Inhibition. J Neurosci 38:1711–1724.

Oberle HM, Ford AN, Dileepkumar D, Czarny J, Apostolides PF (2022) Synaptic mechanisms of top-down control in the non-lemniscal inferior colliculus. eLife 10:e72730.

O’Connor DH, Peron SP, Huber D, Svoboda K (2010) Neural activity in barrel cortex underlying vibrissa-based object localization in mice. Neuron 67:1048–1061.

Pachitariu M, Stringer C, Dipoppa M, Schröder S, Rossi LF, Dalgleish H, Carandini M, Harris KD (2016) Suite2p: beyond 10,000 neurons with standard two-photon microscopy. Neuroscience. Available at: http://biorxiv.org/lookup/doi/10.1101/061507 [Accessed June 11, 2021].

Park J, Phillips JW, Guo J-Z, Martin KA, Hantman AW, Dudman JT (2022) Motor cortical output for skilled forelimb movement is selectively distributed across projection neuron classes. Sci Adv 8:eabj5167.

Payeur A, Guerguiev J, Zenke F, Richards BA, Naud R (2021) Burst-dependent synaptic plasticity can coordinate learning in hierarchical circuits. Nat Neurosci.

Penzo MA, Peña JL (2009) Endocannabinoid-mediated long-term depression in the avian midbrain expressed presynaptically and postsynaptically. J Neurosci Off J Soc Neurosci 29:4131–4139.

Pérez-Garci E, Gassmann M, Bettler B, Larkum ME (2006) The GABAB1b isoform mediates long-lasting inhibition of dendritic Ca2+ spikes in layer 5 somatosensory pyramidal neurons. Neuron 50:603–616.

Peron SP, Freeman J, Iyer V, Guo C, Svoboda K (2015) A Cellular Resolution Map of Barrel Cortex Activity during Tactile Behavior. Neuron 86:783–799.

Peters AJ, Lee J, Hedrick NG, O’Neil K, Komiyama T (2017) Reorganization of corticospinal output during motor learning. Nat Neurosci 20:1133–1141.

Pnevmatikakis EA, Soudry D, Gao Y, Machado TA, Merel J, Pfau D, Reardon T, Mu Y, Lacefield C, Yang W, Ahrens M, Bruno R, Jessell TM, Peterka DS, Yuste R, Paninski L (2016) Simultaneous Denoising, Deconvolution, and Demixing of Calcium Imaging Data. Neuron 89:285–299.

Polley DB, Steinberg EE, Merzenich MM (2006) Perceptual learning directs auditory cortical map reorganization through top-down influences. J Neurosci Off J Soc Neurosci 26:4970–4982.

Quass GL, Rogalla MM, Ford AN, Apostolides PF (2023) Mixed representations of sound and action in the auditory midbrain. Neuroscience. Available at: http://biorxiv.org/lookup/doi/10.1101/2023.09.19.558449 [Accessed September 20, 2023].

Raman IM, Tong G, Jahr CE (1996) Beta-adrenergic regulation of synaptic NMDA receptors by cAMP-dependent protein kinase. Neuron 16:415–421.

Ranganathan GN, Apostolides PF, Harnett MT, Xu N-L, Druckmann S, Magee JC (2018) Active dendritic integration and mixed neocortical network representations during an adaptive sensing behavior. Nat Neurosci 21:1583–1590.

Raymond JL, Medina JF (2018) Computational Principles of Supervised Learning in the Cerebellum. Annu Rev Neurosci 41:233–253.

Rothschild G, Nelken I, Mizrahi A (2010) Functional organization and population dynamics in the mouse primary auditory cortex. Nat Neurosci 13:353–360.

Ruediger S, Scanziani M (2020) Learning speed and detection sensitivity controlled by distinct cortico-fugal neurons in visual cortex. eLife 9.

Schmitt TTX, Andrea KMA, Wadle SL, Hirtz JJ (2023) Distinct topographic organization and network activity patterns of corticocollicular neurons within layer 5 auditory cortex. Front Neural Circuits 17:1210057.

Schneider DM, Nelson A, Mooney R (2014) A synaptic and circuit basis for corollary discharge in the auditory cortex. Nature 513:189–194.

Schoenfeld G, Kollmorgen S, Lewis C, Bethge P, Reuss AM, Aguzzi A, Mante V, Helmchen F (2021) Dendritic integration of sensory and reward information facilitates learning. Neuroscience. Available at: http://biorxiv.org/lookup/doi/10.1101/2021.12.28.474360 [Accessed June 29, 2022].

Schofield BR (2009) Projections to the inferior colliculus from layer VI cells of auditory cortex. Neuroscience 159:246–258.

Shaheen LA, Slee SJ, David SV (2021) Task Engagement Improves Neural Discriminability in the Auditory Midbrain of the Marmoset Monkey. J Neurosci Off J Soc Neurosci 41:284–297.

Sherman SM, Usrey WM (2021) Cortical control of behavior and attention from an evolutionary perspective. Neuron:S0896-6273(21)00462–1.

Slater BJ, Willis AM, Llano DA (2013) Evidence for layer-specific differences in auditory corticocollicular neurons. Neuroscience 229:144–154.

Solyga M, Barkat TR (2021) Emergence and function of cortical offset responses in sound termination detection. eLife 10:e72240.

Stringer C, Pachitariu M, Steinmetz N, Reddy CB, Carandini M, Harris KD (2019) Spontaneous behaviors drive multidimensional, brainwide activity. Science 364:255.

Suga N, Xiao Z, Ma X, Ji W (2002) Plasticity and corticofugal modulation for hearing in adult animals. Neuron 36:9–18.

Sutton RS, Barto AG (2018) Reinforcement learning: an introduction, Second edition. Cambridge, Massachusetts: The MIT Press.

Takahashi H, Magee JC (2009) Pathway interactions and synaptic plasticity in the dendritic tuft regions of CA1 pyramidal neurons. Neuron 62:102–111.

Takahashi N, Ebner C, Sigl-Glöckner J, Moberg S, Nierwetberg S, Larkum ME (2020) Active dendritic currents gate descending cortical outputs in perception. Nat Neurosci 23:1277–1285.

Takahashi N, Oertner TG, Hegemann P, Larkum ME (2016) Active cortical dendrites modulate perception. Science 354:1587–1590.

Takamiya S, Shiotani K, Ohnuki T, Osako Y, Tanisumi Y, Yuki S, Manabe H, Hirokawa J, Sakurai Y (2022) Auditory Cortex Neurons Show Task-Related and Learning-Dependent Selectivity toward Sensory Input and Reward during the Learning Process of an Associative Memory Task. eNeuro 9:ENEURO.0046-22.2022.

Takesian AE, Bogart LJ, Lichtman JW, Hensch TK (2018) Inhibitory circuit gating of auditory critical-period plasticity. Nat Neurosci 21:218–227.

Tang L, Higley MJ (2020) Layer 5 Circuits in V1 Differentially Control Visuomotor Behavior. Neuron 105:346–354.e5.

Tanner WP, Swets JA (1954) A decision-making theory of visual detection. Psychol Rev 61:401–409.

Tervo DGR, Hwang B-Y, Viswanathan S, Gaj T, Lavzin M, Ritola KD, Lindo S, Michael S, Kuleshova E, Ojala D, Huang C-C, Gerfen CR, Schiller J, Dudman JT, Hantman AW, Looger LL, Schaffer DV, Karpova AY (2016) A Designer AAV Variant Permits Efficient Retrograde Access to Projection Neurons. Neuron 92:372–382.

Trautmann EM et al. (2021) Dendritic calcium signals in rhesus macaque motor cortex drive an optical brain-computer interface. Nat Commun 12:3689.

Tsodyks MV, Markram H (1997) The neural code between neocortical pyramidal neurons depends on neurotransmitter release probability. Proc Natl Acad Sci 94:719–723.

Usrey WM, Sherman SM (2019) Corticofugal circuits: Communication lines from the cortex to the rest of the brain. J Comp Neurol 527:640–650.

Williams SR, Atkinson SE (2007) Pathway-specific use-dependent dynamics of excitatory synaptic transmission in rat intracortical circuits: Synaptic dynamics in cortical circuits. J Physiol 585:759–777.

Williams SR, Fletcher LN (2019) A Dendritic Substrate for the Cholinergic Control of Neocortical Output Neurons. Neuron 101:486–499.e4.

Williamson RS, Polley DB (2019) Parallel pathways for sound processing and functional connectivity among layer 5 and 6 auditory corticofugal neurons. eLife 8:e42974.

Winer JA, Larue DT, Diehl JJ, Hefti BJ (1998) Auditory cortical projections to the cat inferior colliculus. J Comp Neurol 400:147–174.

Xu N, Harnett MT, Williams SR, Huber D, O’Connor DH, Svoboda K, Magee JC (2012a) Nonlinear dendritic integration of sensory and motor input during an active sensing task. Nature 492:247– 251.

Xu W, Morishita W, Buckmaster PS, Pang ZP, Malenka RC, Südhof TC (2012b) Distinct neuronal coding schemes in memory revealed by selective erasure of fast synchronous synaptic transmission. Neuron 73:990–1001.

Yan W, Suga N (1998) Corticofugal modulation of the midbrain frequency map in the bat auditory system. Nat Neurosci 1:54–58.

Yang Y, Lee J, Kim G (2020) Integration of locomotion and auditory signals in the mouse inferior colliculus. eLife 9:e52228.

Yu L, Hu J, Shi C, Zhou L, Tian M, Zhang J, Xu J (2021) The causal role of auditory cortex in auditory working memory. eLife 10:e64457.

Yudintsev G, Asilador AR, Sons S, Vaithiyalingam Chandra Sekaran N, Coppinger M, Nair K, Prasad M, Xiao G, Ibrahim BA, Shinagawa Y, Llano DA (2021) Evidence for Layer-Specific Connectional Heterogeneity in the Mouse Auditory Corticocollicular System. J Neurosci Off J Soc Neurosci 41:9906–9918.

Zhan XJ, Cox CL, Sherman SM (2000) Dendritic depolarization efficiently attenuates low-threshold calcium spikes in thalamic relay cells. J Neurosci Off J Soc Neurosci 20:3909–3914.

Zhang LI, Bao S, Merzenich MM (2001) Persistent and specific influences of early acoustic environments on primary auditory cortex. Nat Neurosci 4:1123–1130.

Znamenskiy P, Zador AM (2013) Corticostriatal neurons in auditory cortex drive decisions during auditory discrimination. Nature 497:482–485.

